# Atg16l1 promotes lung transplant tolerance by regulating glycolysis in macrophages

**DOI:** 10.1101/2025.10.02.679826

**Authors:** Marlene Cano, Fuyi Liao, Dequan Zhou, Catherine Chen, Zhiyi Liu, Jihong Zhu, Cory Bernadt, Ricky Ebenezer, Victoria Davis, Katy N. Pugh, Yan Tao, Laneshia K. Tague, Howard J. Huang, Derek E. Byers, Ramsey R. Hachem, Steven L. Brody, Alexander S. Krupnick, Daniel Kreisel, Andrew E. Gelman

## Abstract

Lung transplant survival is limited by the development of chronic lung allograft dysfunction (CLAD), a type of graft rejection that lacks effective treatments. Autophagy plays a crucial role in maintaining cellular homeostasis. In a single-nucleotide polymorphism screen, we found that lung recipients with two copies of a common hypofunctional genetic variant of autophagy-related 16-like 1 rs2241880 (*ATG16L1^T300A/T300A^*), known to deplete this protein from macrophages, were more likely to develop early CLAD. To understand this, we used a mouse orthotopic lung transplant model. Recipients encoding myeloid cell-specific deletion of *Atg16l1* (*Atg16l1^Δ/Δ^*) or who harbor an engineered orthologous mutation (*Atg16l1^T316A/T316A^*) showed similar susceptibility to CLAD. Transcript profiling and mitochondrial tracking studies indicated that increased mitochondrial damage and decreased autophagic removal of mitochondria in Atg16l1-deficient macrophages were associated with heightened activation of the hypoxia-inducible factor 1α (Hif1α) pathway and accumulation of glycolytic transcripts. Metabolic analysis revealed reduced oxidative phosphorylation, increased glycolytic activity, and higher IL-1β expression in Atg16l1-deficient macrophages. Notably, the development of CLAD in *Atg16l1^Δ/Δ^* lung recipients could be significantly prevented by additionally deleting Hif1α in myeloid cells or by treating with the glycolysis inhibitor 2-deoxyglucose. Our results show how a common autophagy-related genetic variant disrupts macrophage metabolism and impairs lung transplant tolerance, pointing toward potential therapeutic strategies to combat CLAD.

## Introduction

Lung transplantation is a life-saving treatment for patients with end-stage pulmonary disease. Unfortunately, overall survival rates are low, with 5-and 10-year survival rates of 57% and 33%, respectively (*26*). Chronic lung allograft dysfunction (CLAD) remains the leading cause of death for lung transplant recipients one year after transplant (*18, 26*). CLAD is characterized by the persistent loss of expiratory graft function and can result from progressive fibrotic obliteration of the small airways, described as Bronchiolitis Obliterans Syndrome (BOS) or restrictive allograft syndrome (RAS), marked by the appearance of interstitial and pleural fibrosis. Despite increased understanding of alloimmune (*12*) and non-alloimmune risk factors (*28, 55, 68*) involved in CLAD development, there is a lack of effective therapies to address this poor outcome.

Previous work has shown that CLAD development is connected to club cell loss or dysfunction, as indicated by reductions of club cell secretory protein (CCSP) (*20, 23*). Club cells are non-ciliated bronchiolar epithelial cells that repair bronchiolar injury through their capacity to self-renew and differentiate into goblet and ciliated cells (*43, 56*). Using a mouse orthotopic lung transplant model (*47*), we recently demonstrated that club cell injury results in immunological responses and allograft tissue injury consistent with CLAD development (*37, 38*). This model utilizes donor lungs on a FVB background that encode three transgenes (3T; H-2^q^), which coordinate the recombination of a Loxp-Stop-Loxp diphtheria toxin A locus by a Cre recombinase under inducible transcriptional control by a reverse tetracycline transactivator protein expressed specifically in club cells (*37*). Transplantation of 3T lungs into immunosuppressed, major histocompatibility-mismatched C57Bl/6 (B6; H-2^b^) recipients, followed by doxycycline ingestion for 2.5 days, leads to transient depletion of club cells in the allografts. After doxycycline uptake, club cell regeneration is poor, resulting in obliterative lesions and marked parenchymal fibrosis resembling severe CLAD pathology. In contrast, doxycycline ingestion by B6 recipients who received 3T lungs on a syngeneic B6 background recovers their club cell compartment and does not develop CLAD (*37, 38*).

Autophagy is an essential homeostatic cellular process that helps eliminate invading pathogens and damaged intracellular macromolecules and organelles by facilitating their degradation. Selective forms of autophagy, such as mitophagy, are associated with various diseases, including Parkinson’s disease, cancer, and infections (*51, 60*). In 2007, a genome-wide association study analysis of nonsynonymous single-nucleotide polymorphisms (SNPs) identified the G substitution at rs2241880 of the autophagy 16-like 1 (*ATG16L1*) gene as a susceptibility variant for Crohn’s disease (*17*). This SNP leads to a threonine-to-alanine (T300A) substitution in ATG16L1. Murthy and colleagues demonstrated that *ATG16l1^T300A/T300A^* macrophages lack this protein, due to the creation of a caspase-3 cleavage site. Under stress conditions that activate caspase 3, ATG16L1 undergoes cleavage, resulting in a deficiency of this protein (*45*). Mice engineered with an orthologous *ATG16L1^T300A/T300A^* mutation (*Atg16l1^T316A/T316A^*) also lack this protein in their macrophages(*45*). *Atg16l1^T316A/T316A^* macrophages, or those that harbor myeloid-specific genetic deletion of *Atg16l1*, exhibit increased inflammatory responses associated with Crohn’s disease progression (*30, 45*).

However, it remains unclear whether Atg16l1 deficiency influences inflammatory responses that affect outcomes after lung transplantation.

Hypoxia-inducible factor 1α (Hif1α) is a labile transcription factor that plays a crucial role in promoting protective responses to oxidative stress and tissue damage. In macrophages, Hif1α stability is enhanced by mitochondrial metabolites generated by LPS stimulation, including superoxide (*66*). Hif1α stabilization, in turn, promotes the metabolic reprogramming of macrophages from oxidative phosphorylation to glycolysis, partly through driving the transcription of genes in the glycolytic pathway (*40*). Hif1α-mediated glycolysis supports the expression of pro-fibrogenic factors, such as IL-1β (*22, 58, 66*). Except for LPS, factors contributing to Hif1α-mediated metabolic alterations remain less defined, and their potential impact on lung transplant tolerance remains unknown (*36*).

Here, we report that lung transplant recipients carrying the GG genotype of *ATG16L1* rs2241880 have an increased risk of early CLAD development. Using a mouse orthotopic lung transplant model, we show that recipients with Atg16l1-deficient macrophages, or that encode *Atg16l1^T316A/T316A^*, develop severe signs of CLAD following only minimal club cell injury. Analysis of Atg16l1-deficient macrophages reveals signs of mitochondrial stress and a defect in mitophagy, accompanied by enhanced Hif1α stabilization and increased glycolytic flux. However, lung recipients with Atg16l1 deficiency in their macrophage compartment are protected from CLAD when Hif1α is genetically deleted or glycolysis is pharmacologically inhibited.

## Results

### Atg16l1 rs2241880 increases the risk of early BOS development

To assess the role of ATG16L1 in lung transplant survival, we genotyped 237 recipients who underwent lung transplantation between 1992 and 2013 at Barnes-Jewish Hospital. In this cohort, 68 (29.7%) lung recipients were homozygous for rs2241880 (GG), 101 (44.1%) were heterozygous for rs2241880 (AG), and 60 (26.2%) did not carry rs2241880 (AA) (**Figure 1A**). In the genotyped recipients, the mean age was 52 years (± 14 years), 141 patients (60%) were male, and 225 patients (95%) were white non-Hispanic, which is representative of the recipients at our lung transplant center (**Table 1 and Supplementary Table 1**). The most common reasons for transplant were interstitial lung disease (ILD, n=104, 43.8%), 5 chronic obstructive pulmonary disease (COPD, n=52, 21.9%), and cystic fibrosis (CF, n=47, 9.8%). Two hundred thirty-two patients (97.9 %) received bilateral lung transplants, and 113 (47.7%) required cardiopulmonary support during transplantation. We also examined infection in genotyped lung recipients (**Supplementary Table 2**) showing that 83 patients (36.2%) grew *Aspergillus*, 63 (27.5%) gram-positive bacteria, of which 46 (20.1%) were *Staphylococcus aureus*, 119 (52%) gram-negative bacteria, of which 53 (23.1%) were *Pseudomonas aeruginosa* and 50 (21.8%) were *Stenotrophomonas maltophilia*. Eighteen (7.9%) recipients had cytomegalovirus (CMV) recovered from their airways, and 137 (59.8%) had CMV in their blood, with only 1 (0.4%) recipient having CMV both in the blood and airways. All transplant patients received acyclovir for CMV prophylaxis and were treated with valganciclovir for CMV detected in the airways or blood. 82 (35.8%) recipients had a community-acquired respiratory virus, of which 5 (2.2%) were Influenza and 16 (6.6%) respiratory syncytial virus. A univariate risk factor analysis for CLAD development identified previously known risk factors, including prior airway infections such as CMV HR 1.91 (0.97-3.7), p=0.055 (**Table 2**). In contrast to previous reports (*14, 24*), we found that a prior episode of CARV was associated with a reduced risk of CLAD, HR 0.61 (0.38-0.96), p=0.033.

**Figure 1.**
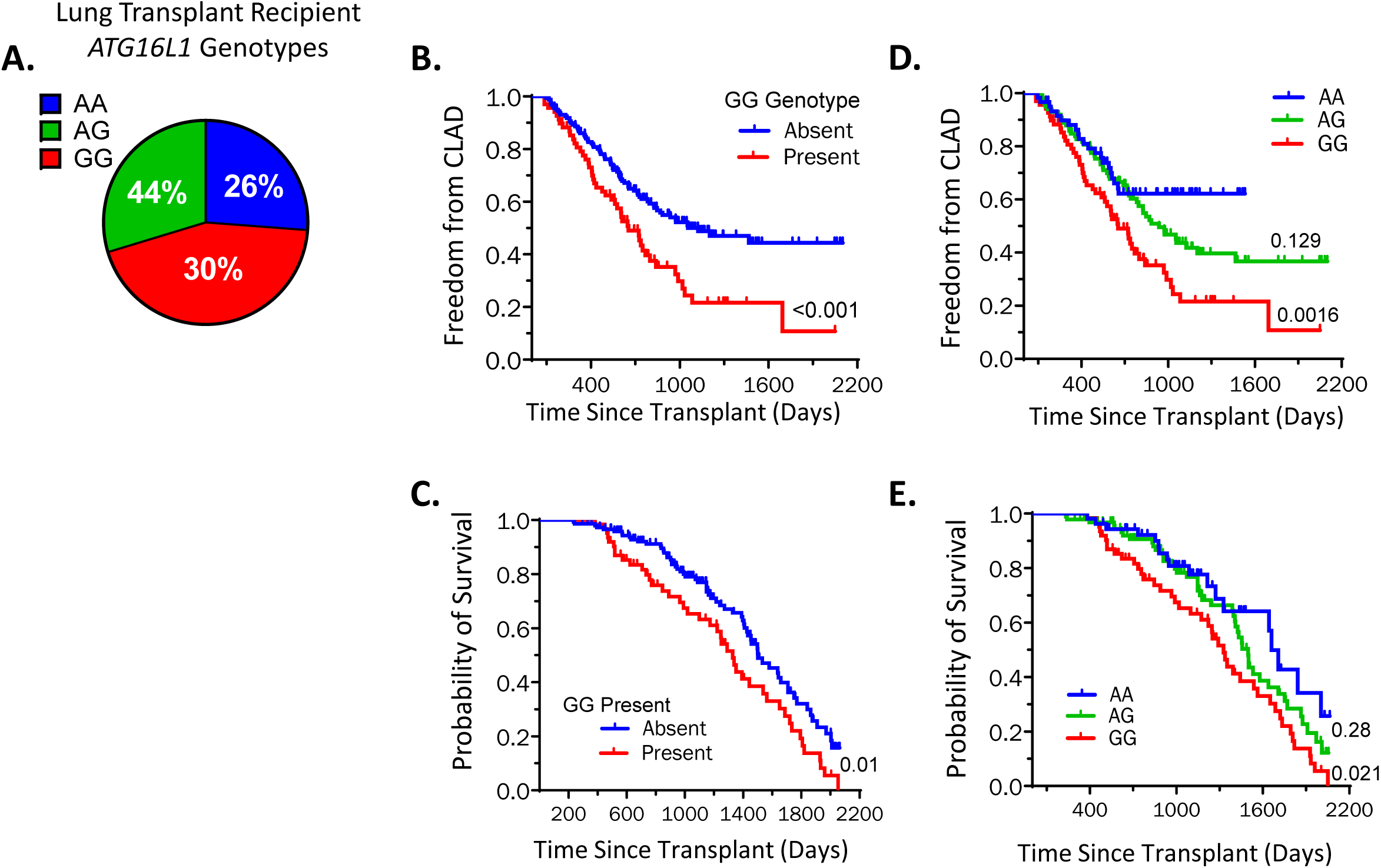
Lung transplant recipients with ATG16L1 (GG) rs2241880 have earlier BOS development. (**A**) Lung transplant recipients genotyped for the *ATG16L1* rs2241880 (n=229). (**B**) Kaplan-Meier curves of the proportion of recipients free from CLAD over time stratified by the presence or absence of GG genotype. Log-rank test for CLAD development, HR 1.84, p=0.0059. (**C**) The probability of transplant survival over time stratified by the presence or absence of GG genotype. Log-rank test for allograft survival, HR 1.65, p=0.01. (**D**) The proportion of recipients free from CLAD over time stratified by AA, AG, or GG genotype. Log-rank test for CLAD development with pairwise comparisons, p=0.0016 between AA and GG, and p=0.129 between AA and AG. (**E**) The probability of transplant survival over time stratified by AA, AG, or GG genotype. Log-rank test with *p<0.05 for AA v GG. Pairwise comparisons between AA and AG did not reach statistical significance with p=0.28. Log-rank test with *P < 0.05 and **p<0.01 and ***p<0.001.

**Table 1.** Demographics and Clinical Characteristics of Lung transplant Recipients in the cohort. Numbers are expressed as n (mean) for continuous and n (%).

| **Demographics and Clinical Characteristics, n=237** | **n (%)** |
| --- | --- |
| Age at time of transplantation, years ± SD | 52 ± 14 |
| Male recipients | 141 (59.5) |
| Recipient race |  |
| White | 225 (94.9) |
| Black | 11 (4.6) |
| Asian | 1 (0.4) |
| Hispanic recipients | 4 (1.7) |
| ABO group |  |
| A | 106 (44.7) |
| B | 22 (9.3) |
| AB | 4 (1.7) |
| O | 105 (44.3) |
| Rh+ Recipients | 200 (84.4) |
| Pretransplant diabetes mellitus | 66 (27.8) |
| Pretransplant pulmonary arterial hypertension | 134 (56.5) |
| CMV positive | 134 (56.5) |
| Pretransplant diagnosis |  |
| A<sub>1</sub>-antitrypsin deficiency emphysema | 12 (5.1) |
| Bronchiectasis unrelated to cystic fibrosis | 8 (3.4) |
| Cystic fibrosis related bronchiectasis | 47 (19.8) |
| Chronic obstructive pulmonary disease (COPD) | 52 (21.9) |
| Emphysema unrelated to COPD | 2 (0.8) |
| Interstitial lung disease (ILD) | 104 (43.8) |
| Lymphangioleiomyomatosis (LAM) | 3 (1.3) |
| Pulmonary arterial hypertension | 3 (1.3) |
| Pulmonary Langerhans cell histiocytosis (PLCH) | 1 (0.4) |
| Sarcoidosis | 5 (2.1) |
| Bilateral lung transplantation | 232 (97.9) |
| Cardiopulmonary bypass | 113 (47.7) |
| ATG16L genotype |  |
| AA | 60 (26.2) |
| AG | 101 (44.1) |
| GG | 68 (29.7) |

**Table 2.** *ATG16l1* rs2241880 GG is an independent risk factor for BOS. Hazard ratio (HR) and 95% confidence interval (CI).

| Risk Factor n=229 | Univariate analysis |  | Multivariate analysis |  |
| --- | --- | --- | --- | --- |
|  | HR (95% CI) | p-value | HR (95% CI) | p-value |
| ATG16L genotype |  |  |  |  |
| AA | 1.00 |  |  |  |
| AG | 1.13 (0.71 – 1.80) | 0.611 |  |  |
| GG | 1.84 (1.14 – 2.98) | 0.012 |  |  |
| ATG16L GG genotype | 1.71 (1.18 – 2.47) | 0.004 | 1.64 (1.12 – 2.38) | 0.011 |
| Previous Gram-positive bacteria | 1.30 (0.87 – 1.94) | 0.203 |  |  |
| *Staphylococcus aureus* | 1.24 (0.79 – 1.94) | 0.348 |  |  |
| Other | 1.53 (0.74 – 3.17) | 0.248 |  |  |
| Previous Gram-negative bacteria | 1.02 (0.72 – 1.46) | 0.906 |  |  |
| *Pseudomonas aeruginosa* | 1.14 (0.74 – 1.75) | 0.560 |  |  |
| *Stenotrophomonas maltophilia* | 0.82 (0.51 – 1.31) | 0.400 |  |  |
| Other | 2.29 (0.92 – 5.72) | 0.076 |  |  |
| Previous cytomegalovirus | 1.28 (0.87 – 1.90) | 0.213 |  |  |
| Blood | 1.22 (0.82 – 1.82) | 0.339 |  |  |
| Airway | 1.91 (0.97 – 3.70) | 0.055 |  |  |
| Blood and airway | 1.74 (0.24 – 12.76) | 0.584 |  |  |
| Previous cytomegalovirus in airway | 1.67 (0.93 – 2.97) | 0.085 | 1.32 (0.73 – 2.40) | 0.355 |
| Previous community-acquired respiratory virus | 0.61 (0.38 – 0.96) | 0.033 | 0.63 (0.39 – 1.00) | 0.048 |
| Influenza | 0.48 (0.07 – 3.45) | 0.467 |  |  |
| Respiratory syncytial virus | 0.49 (0.16 – 1.56) | 0.227 |  |  |
| Adenovirus | 0.000 | 0.979 |  |  |
| Other | 0.65 (0.39 – 1.07) | 0.091 |  |  |
| Previous *Aspergillus* | 1.26 (0.88 – 1.82) | 0.209 |  |  |
| *Aspergillus fumigatus* | 1.26 (0.70 – 2.28) | 0.436 |  |  |
| *Aspergillus niger* | 1.15 (0.58 – 2.29) | 0.697 |  |  |
| *Aspergillus flavus* | 1.70 (0.90 – 3.21) | 0.101 |  |  |
| Other single species | 1.22 (0.63 – 2.35) | 0.563 |  |  |
| Multiple single species | 0.91 (0.33 – 2.50) | 0.860 |  |  |

Strikingly, a significantly increased risk of developing CLAD was observed in lung transplant recipients homozygous for GG rs2241880, HR 1.71 (1.18-2.47), p=0.004. In a multivariate analysis, GG carriers remained at considerable risk for CLAD, HR 1.64 (1.12-2.38), p=0.011, after accounting for CARV and CMV infections. We then assessed whether carrying rs2241880 GG affected freedom from CLAD (**Figure 1B**). While the median time of freedom from CLAD was 1,124 days in lung recipients without GG, it sharply decreased to 654 days in GG carriers. Allograft failure was defined as patients who died from allograft-related dysfunction, including CLAD. Lung recipients with GG developed allograft failure significantly earlier than those without GG (**Figure 1C**), with non-GG recipients reaching a median survival of 1501 days and GG carriers only 1327 days. To determine if carrying one G susceptibility allele affected allograft survival, we divided the non-GG group into AA and AG recipients. Lung recipients with one G allele showed an intermediate time to CLAD (**Figure 1D**) and an intermediate time to allograft failure (**Figure 1E**), trending toward but not reaching statistical significance compared to AA recipients. Carrying *ATG5* variants has been linked to chronic inflammation and airway remodeling (*57*). We genotyped lung recipients for *ATG5* SNPs rs12201458, rs573775, and rs510432; however, carrying these SNPs did not impact freedom from CLAD development, nor was it associated with allograft failure (**Supplementary Figure 1**).

### Atg16l1 expression in recipient-derived macrophages prevents CLAD

To study the function of Atg16l1 in graft survival, we utilized a mouse orthotopic left lung transplant model that develops severe CLAD following transient club cell loss (*37*).

Given that previous work has shown that the Atg16l1 T300A mutation leads to the depletion of Atg16l1 protein in macrophages (*45*), we generated *Lyz2^Cre^Atg16l1^loxP/loxP^* (*Atg16l1^Δ/Δ^*) mice, which have a targeted ablation of the *Atg16l1* gene in myeloid cells. We then transplanted 3T donor lungs into immunosuppressed *Atg16l1^Δ/Δ^* and wildtype control *Atg16l1^loxP/loxP^* (*Atg16l1^fl/fl^*) recipients (**Figure 2A)**. On post-operative day (POD) 7, lung recipients ingested doxycycline for 18 hours, a feeding period that is insufficient to trigger severe CLAD development in wild-type recipients (*37*). Allografts were examined on POD 16 (**Figure 2B**). *Atg16l1^Δ/Δ^* lung recipients had more significant histological evidence of obliterative fibrotic airway lesions and collagenous alveolar exudates when compared to wildtype recipients. Blinded modified Ashcroft scoring indicated more allograft fibrotic injury in *Atg16l1^Δ/Δ^* compared to control recipients (**Figure 2C).** Hydroxyproline accumulation, a biomarker of new lung collagen synthesis and fibrosis, was higher in allografts of *Atg16l1^Δ/Δ^* compared to control recipients (33.9μg/mL vs 21.55 μg/mL, p=0.0053) (**Figure 2D**). We have previously demonstrated that poor club cell regeneration following transient allograft club cell depletion is associated with CLAD development (*37*). Compared to wildtype *Atg16l1^fl/fl^* hosts, *Atg16l1^Δ/Δ^* recipients had significantly fewer lung graft total and proliferating club cells (**Figure 2E, F**). Prior work has shown that macrophages in *Atg16l1^T316A/T316A^* mice also lack Atg16l1 protein (*45*). Similar to *Atg16l1^Δ/Δ^* hosts, *Atg16l1^T316A/T316A^* lung recipients developed severe CLAD (**Supplementary Figure 2**).

**Figure 2.**
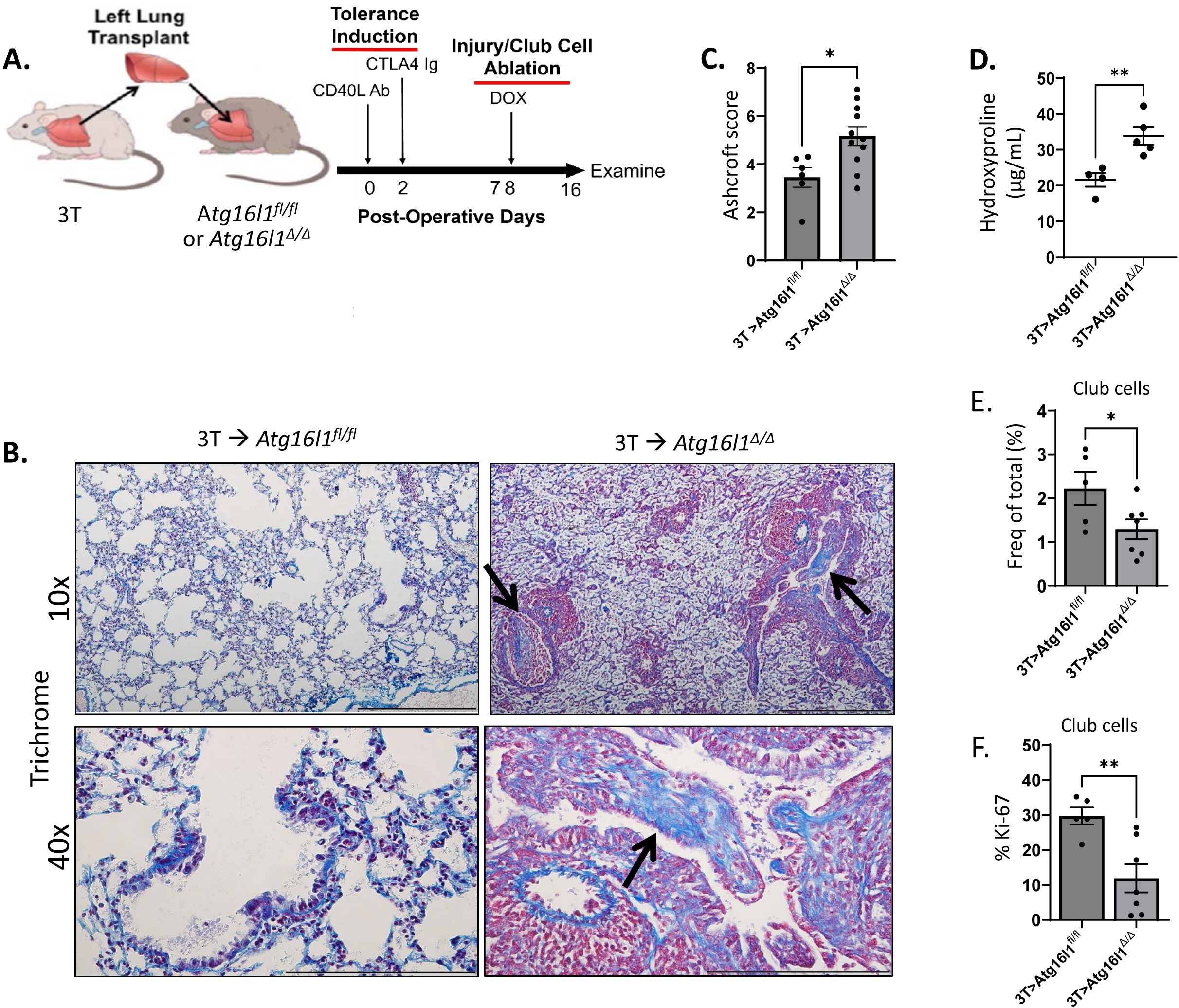
Atg16l1 expression in recipient-derived macrophages prevents CLAD development (**A**) 3T left lungs were orthotopically transplanted into *Atg16l1^Δ/Δ^* and control *Atg16l1^fl/fl^* B6 recipients and treated with CD40L Ab (POD 0) and CTLA4 Ig (POD 2) to establish allograft tolerance. Doxycycline (DOX) was administered in chow and water for 30 hours to induce club cell ablation on post-operative day (POD) 7, and graft survival, and epithelial injury were analyzed on POD 16. (**B**) Representative allograft Gomori Trichrome staining. Images shown are representative of n≥7/group. Arrows denote fibrotic lesions. (**C**) Randomly selected trichrome-stained images of lung grafts were scored for fibrosis and graded blindly using a modified Ashcroft scale. Allografts were evaluated for (**D**) hydroxyproline content (n≥4/group), (**E**) total club cell (n≥5/group), and (**F**) Ki-67^+^ club cell (n≥5/group) recovery relative to the CD45^-^ cell compartment. Data are represented as mean ± SEM. Two-sided Mann-Whitney U test (**C-F**). *P < 0.05; **P < 0.01. Scale bars: 500 μm for 10x, 200 μm for 40x.

### Reduced mitophagy in Atg16l1-deficient macrophages

Our previous work has shown that recipient monocytes give rise to monocyte-derived alveolar macrophages (Mo-AM) that promote CLAD development (*38*). We performed bulk RNA sequencing on FACS-sorted Mo-AM from lung allografts that had been transplanted into *Atg16l1^Δ/Δ^* or *Atg16l1^fl/fl^* recipients **(Figure 3A**). Gene ontology analysis revealed the downregulation of several pathways involved in mitochondrial respiration in *Atg16l1^Δ/Δ^* Mo-AMs. This finding was linked to a sharp reduction of nuclear-encoded electron transport complex (ETC) transcripts in *Atg16l1^Δ/Δ^* relative to wildtype Mo-AMs (**Figure 3B**). Because these observations suggested mitochondrial injury, we evaluated mitochondrial mass, membrane potential, and mitochondrial-specific reactive oxygen species (ROS) production in *Atg16l1^Δ/Δ^* bone marrow monocyte-generated macrophages (BMM) (**Figure 4A**). When compared to *Atg16l1^fl/fl^* BMM, mitochondrial mass, membrane potential, and ROS production were higher in *Atg16l1^Δ/Δ^* BMM. The high mitochondrial mass and elevated ROS production suggested a defect in mitophagy, the selective autophagic removal of damaged mitochondria (*8*). To examine whether this was the case, we crossed *Atg16l1^T316A/T316A^* mice with mitophagy reporter mice that express a mitochondrially targeted Kiema protein (mt-Keima). In these mice, a pH-sensitive, dual-excitation fluorescent mt-Keima protein distinguishes mitochondria residing in the cytoplasm from those localized within lysosomes (*64*). Compared to wildtype macrophages, there was significantly less mitochondrial flux into lysosomes of *Atg16l1^T316A/T316A^* macrophages (**Figures 4B, C**). Atg16l1 associates with mitochondria in embryonic fibroblasts to promote mitophagy, but whether it does so in macrophages remains unclear (*21*). In wildtype BMM, we observed colocalization of Atg16l1 and mitochondria (**Figure 4D, Supplementary Movie 1**).

**Figure 3.**
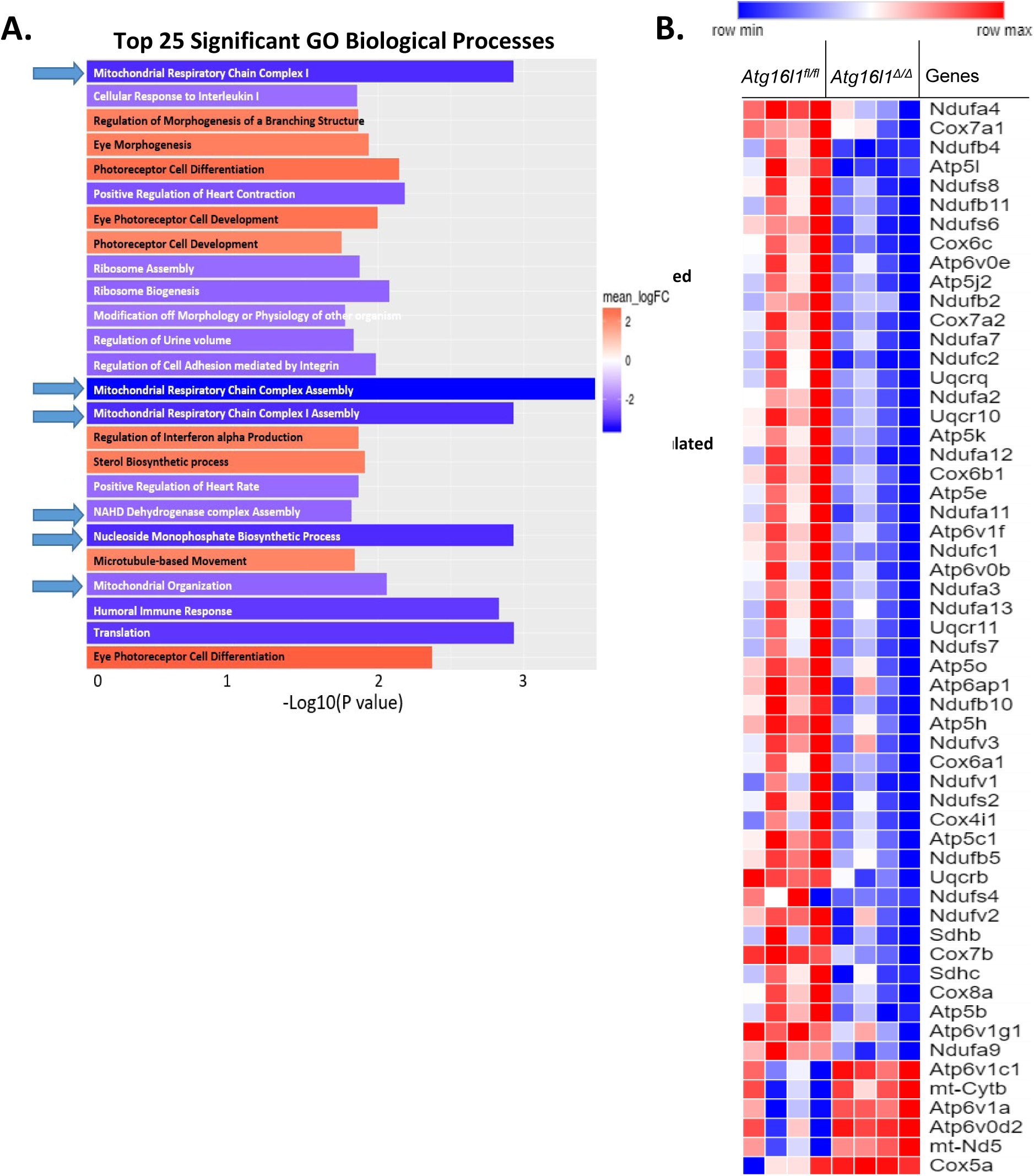
Mitochondrial electron transport complex (ETC) transcripts are downregulated in *Atg16l1^Δ/Δ^* alveolar macrophages (AM). Bulk RNA sequencing of FACS-sorted AM from *Atg16l1^Δ/Δ^* and *Atg16l1^fl/fl^* mice (**A**) Top significant GO biological process pathways and (**B**) heatmap of ETC transcripts that had changes in accumulation of p<0.05 (n=4 per group). Arrows in (**A**) denote pathways related to mitochondrial function and metabolism.

**Figure 4.**
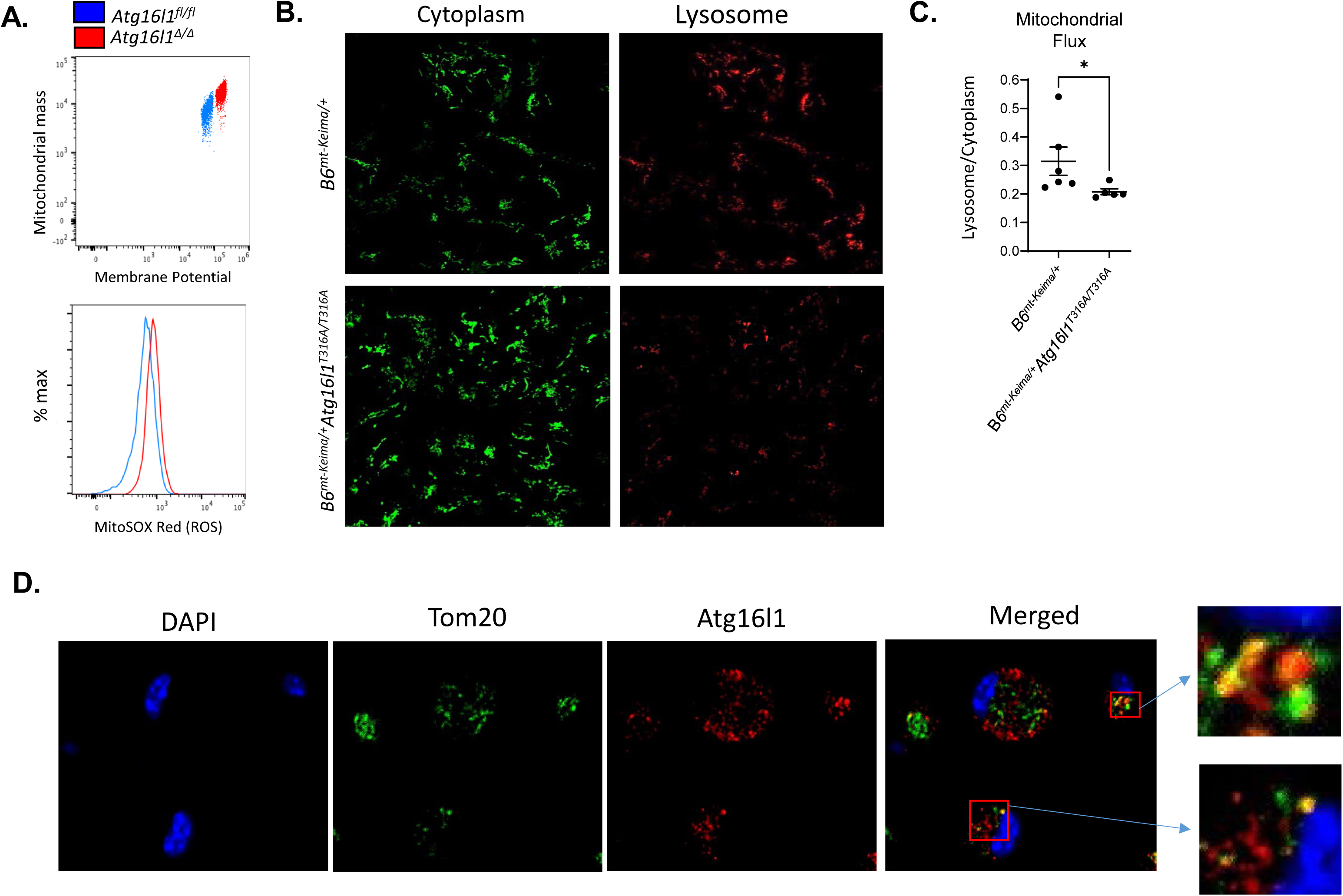
**Increased ROS production and reduced mitophagy in *Atg16l1^T316A/T316A^*** BMM (**A**) A representative flow cytometric analysis (n=3) of mitochondrial mass versus membrane potential and mitochondrial ROS production in resting *Atg16l1^Δ/Δ^* and *Atg16l1^fl/fl^* BMMs. (**B**) Assessment of mitophagic flux using mt-keima reporter mice. Confocal images of *B6^mt-Keima/+^Atg16l1^Δ/Δ^* and *B6^mt-Keima/+^Atg16l1^fl/fl^* BMMs were analyzed for cytoplasmic (E_m_ 488nm (pH ∼ 7)) and lysosome-localized (E_m_ 568nm (pH ∼ 4)) mitochondria. (**C**) A plot of the ratio of lysosomal to cytoplasmic localized mitochondria as measured by mean fluorescence intensity of mitochondria within each compartment (n>3/group). (**D**) Confocal imaging of the colocalization of Atg16l1 and mitochondria in BMMs. The images shown are representative of mitochondria transmembrane protein Tom20 and Atg16l1 co-staining (n = 3). (**A-C**) Pearson’s correlation coefficient between Tom20 and Atg16l1 R=0.34. Insets show co-localization between mitochondria and Atg16l1 two-sided Mann-Whitney U test. *P < 0.05.

### Elevated glycolysis and Hif1α-stabilization in Atg16l1-deficient macrophages

Mitochondrial ETC dysfunction is linked to increased glycolysis (*4*), suggesting that Atg16l1 deficiency may enhance glucose metabolism. Extracellular proton flux analysis of *Atg16l1^Δ/Δ^* BMM demonstrated elevated baseline glycolysis relative to *Atg16l1^fl/fl^* BMM (**Figure 5A**). *Atg16l1^Δ/Δ^* macrophages exhibited increases in glycolytic capacity and reserve, measures of energetic demand, and maximal glycolytic-mediated ATP production, respectively. To validate these observations, we assessed lactate release and radiometric glucose flux in *Atg16l1^Δ/Δ^* (**Figures 5B, C**) and *Atg16l1^T316A/T316A^* BMMs (**Supplemental Figure 3**). In line with extracellular flux measurements, *Atg16l1^Δ/Δ^* and *Atg16l1^T316A/T316A^* BMM produced more lactate and displayed higher rates of glycolytic flux relative to *Atg16l1^fl/fl^* BMM. Conversely, mitochondrial-mediated ATP generation, along with basal and maximal respiration, was significantly reduced in *Atg16l1^Δ/Δ^* macrophages as measured by oxygen consumption rate (OCR) (**Figure 5D**).

**Figure 5.**
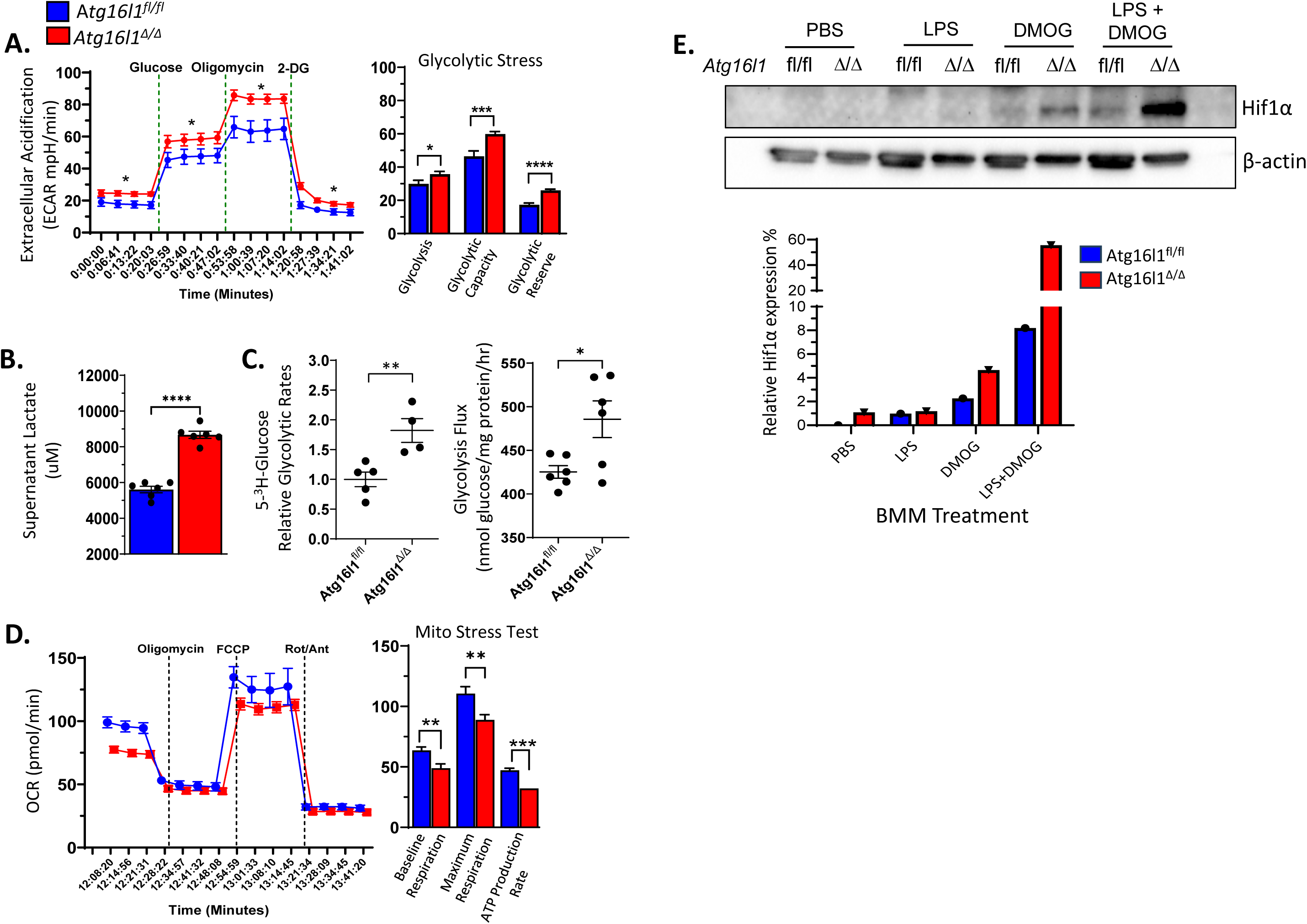
Reduced oxidative phosphorylation and increased glycolysis in *Atg16l1^Δ/Δ^* BMM. (**A**) Extracellular acidification rate (ECAR) analysis conducted on *Atg16l1^Δ/Δ^* and wildtype *Atg16l1^fl/fl^* BMMs. Data shown is a representative result from three independent experiments. (**B**) Baseline supernatant lactate concentrations of *Atg16l1^Δ/Δ^* and control *Atg16l1^fl/fl^* BMMs (n=6/group). (**C**) Baseline radiometric glycolytic flux for indicated BMMs following incubation with D-[5-^3^H(N)] labeled glucose. (**D**) Changes in oxygen consumption rate (OCR) in response to treatment with indicated electron transport complex inhibitors from a mitochondrial stress test. (**E**) Protein expression of Hif1α relative to β-actin in *Atg16l1^Δ/Δ^* and control *Atg16l1^fl/fl^* BMMs (representative of 2 independent experiments). ECAR and OCR plots show a representative result from n≥3 experiments, 5-7 replicates per experiment, where a Mann-Whitney U-test was used to evaluate significance; *P < 0.05; **P < 0.01, ****P < 0.0001.

Glycolytic expression is controlled by transcription factor Hif1α, and LPS stimulation of BMM promotes the stabilization of Hif1α (*5, 22, 32, 46, 58, 66*).

Dimethyloxalylglycine (DMOG) antagonizes the degradation of Hif1α, which allows for the visualization of this protein under normal oxygen tension. We analyzed Hif1α protein levels in cultured BMMs following treatment with LPS in combination with DMOG treatment. When compared to *Atg16l1^fl/fl^* BMM, *Atg16l1^Δ/Δ^* BMM had substantially more Hif1α protein accumulation following LPS plus DMOG stimulation relative to DMOG stimulation alone (**Figure 5E**). Additionally, bulk-RNA sequencing of flow cytometric-sorted *Atg16l1^Δ/Δ^* Mo-AM, monocyte-derived interstitial macrophages, and cultured BMM all showed enrichment for Hif1α transcript levels relative to the same cells from *Atg16l1^fl/fl^* mice (**Supplemental Figure 4A, B**). Additionally, glycolysis-related enzyme transcript levels were higher in *Atg16l1^Δ/Δ^* when compared to *Atg16l1^fl/fl^* BMM (**Supplemental Figure 4C)**. Hif1α drives IL-1β gene transcription (*11, 66*). Compared to *Atg16l1^fl/fl^* BMMs, *Atg16l1^Δ/Δ^* BMMs showed significantly more spontaneous and LPS-mediated IL-1β production (**Supplemental Figure 4D)**.

### Antagonizing glycolysis prevents CLAD

To determine if augmented glycolysis in Atg16l1-deficient macrophages is Hif1α-dependent, we generated *Lyz2^Cre^ Atg16l1^fl/fl^ Hif1α^fl/fl^* (*Atg16l1^Δ/Δ^ Hif1α ^Δ/Δ^*) mice.

Consistent with previous observations on the critical Hif1α transcriptional activity in promoting glycolysis (*40*), levels of glycolysis-associated gene transcripts *Pgk1, Pdk1, Ldha, Gdpd4*, and *Hk2* were lowered by *Hif1α* gene deletion in LPS-stimulated *Atg16l1^Δ/Δ^* macrophages (**Supplemental Figure 5**). Additionally, baseline lactate accumulation and IL-1β production were sharply reduced in *Atg16l1^Δ/Δ^ Hif1α ^Δ/Δ^* relative to *Atg16l1^Δ/Δ^* BMMs. However, LPS-stimulated IL-1β production was not wholly eliminated from *Atg16l1^Δ/Δ^ Hif1α ^Δ/Δ^* BMMs, indicating the presence of other pathways that can control the expression of this cytokine.

To evaluate the role of Hif1α in CLAD development in *Atg16l1^Δ/Δ^* lung recipients, we transplanted 3T lungs into allogeneic *Atg16l1^Δ/Δ^ Hif1α^Δ/Δ^* recipients. There was less histological evidence of obliterative airway lesions in allografts after transplantation into *Atg16l1^Δ/Δ^ Hif1α^Δ/Δ^* compared to *Atg16l1^Δ/Δ^* recipients (**Figures 6A, B**). Consistent with these observations, *Hif1α* gene deletion led to a 1.6-fold increase in recovery of club cell numbers (**Figure 6C**). We next determined if Hif1α regulated IL-1β expression in intragraft macrophages. We noted a significant reduction in graft-resident IL-1β^+^ alveolar macrophages (**Figure 6D**) and IL-1β^+^ CD11c^+^ cells (**Figure 6E**) in allografts after transplantation into *Atg16l1^Δ/Δ^ Hif1α^Δ/Δ^* compared to *Atg16l1^Δ/Δ^* recipients. Plasma lactate levels, a surrogate indicator of glycolysis, were also reduced in *Atg16l1^Δ/Δ^ Hif1α^Δ/Δ^* lung recipients (**Figure 6F**). To further dissect the contribution of Hif1α to allograft survival, we transplanted 3T lungs into *Atg16l1^Δ/Δ^* recipients where only one *Hif1α* allele was deleted (*Atg16l1^Δ/Δ^ Hif1α^Δ/+^*) (**Supplementary Figure 6**). Allograft injury in *Atg16l1^Δ/Δ^ Hif1α^Δ/+^* recipients was less relative to that of *Atg16l1^Δ/Δ^* recipients but comparable to *Atg16l1^Δ/Δ^ Hif1α^Δ/Δ^* recipients (**Supplemental Figure 6 versus Figures 6A and B**). However, lactate accumulation and IL-1β expression in the supernatant of *Atg16l1^Δ/Δ^ Hif1α^Δ/+^* BMMs were reduced compared to *Atg16l1^Δ/Δ^* but remained higher than in *Atg16l1^Δ/Δ^ Hif1α^Δ/Δ^* BMMs (**Supplementary Figure 7**).

**Figure 6.**
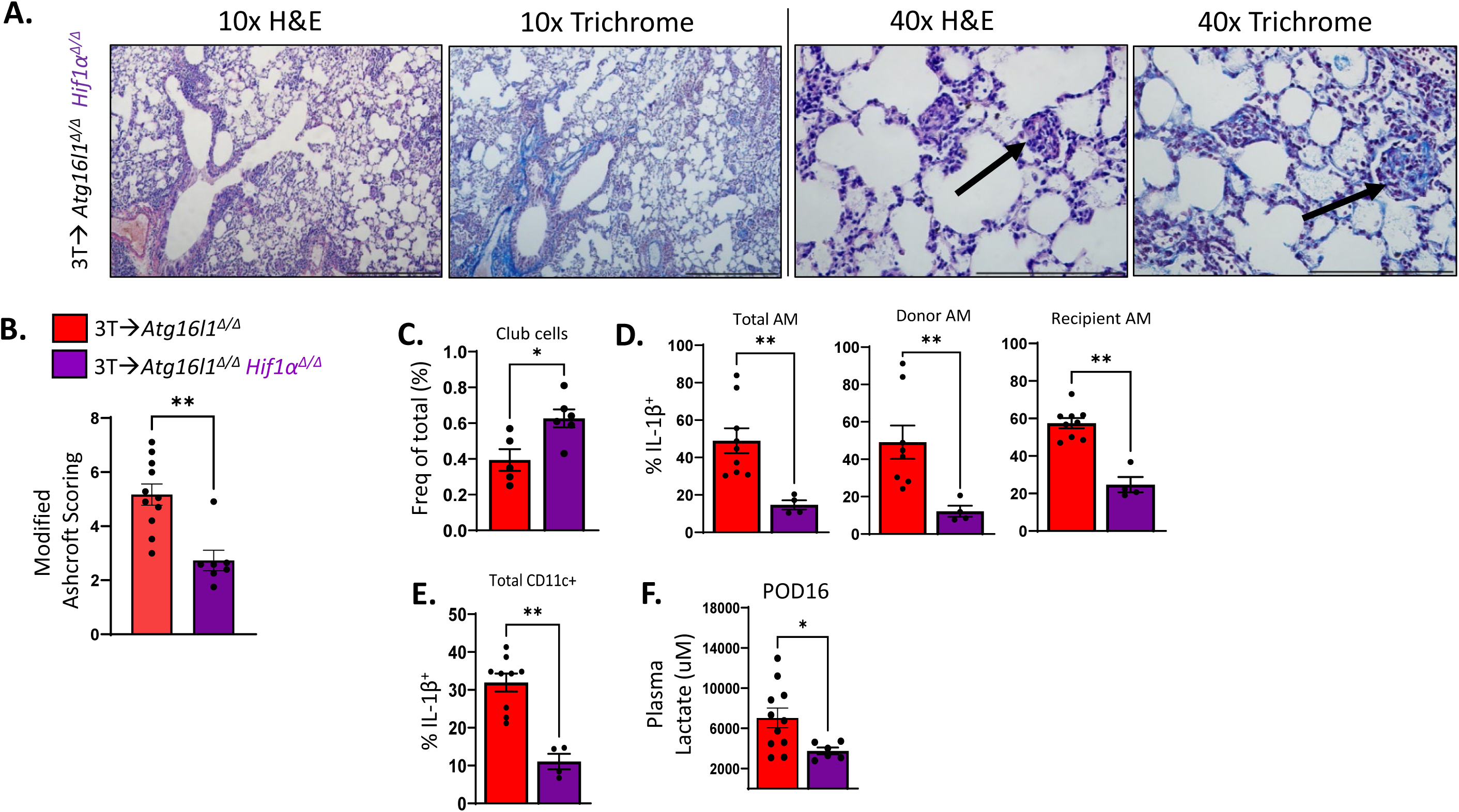
CLAD development in Atg16l1^Δ/Δ^ lung recipients is Hif1α dependent 3T FVB lung allografts of *Atg16l1^Δ/Δ^* and *Atg16l1^Δ/Δ^ Hif1α^Δ/Δ^* recipients analyzed for evidence of fibrosis on POD16. (**A**) Representative images of allograft H&E and trichrome staining (n≥6/group). (**B**) Allograft tissue blindly graded using modified Ashcroft scoring. POD 16 allografts assessed for (**C**) club cell abundance relative to CD45^-^ cells, (**D**) intracellular AM IL-1β expression, and (**E**) the percent abundance of IL-1β^+^ CD11c^+^ cells. (**F**) Lung recipient plasma was collected on POD 16 and evaluated for lactate levels. Data in **B-F** are represented as mean ± SEM. Mann U Whitney T-test (**C-F**). *P < 0.05; **P < 0.01, ***P < 0.001, ****P < 0.0001. Scale bars: 500 μm for 10x, 200 μm for 40x.

We next inquired whether pharmacologically inhibiting glycolysis could prevent CLAD development in *Atg16l1^Δ/Δ^* recipients. The glycolytic inhibitor 2-deoxyglucose (2-DG) has been demonstrated to reduce Hif1α-dependent IL-1β production (*66*) and has been evaluated in clinical trials for cancer treatment and Covid-19 infection (*2, 41, 48, 63*). 2-DG treatment inhibited lactate and IL-1β production in LPS-stimulated *Atg16l1^Δ/Δ^*

BMMs in a dose-dependent manner (**Figures 7A, B**). To determine if blocking glycolysis prevents CLAD pathogenesis in *Atg16l1^Δ/Δ^* lung recipients, we administered 2-DG or vehicle control (PBS) daily from POD 7 to 16. 2-DG-treated *Atg16l1^Δ/Δ^* lung recipients exhibited less histological evidence of allograft injury compared to vehicle-treated *Atg16l1^Δ/Δ^* lung recipients (**Figure 7C, D**). 2-DG treatment increased allograft club cell levels by approximately 3.4-fold relative to vehicle treatment (**Figure 7E**), suggesting that inhibiting glycolysis promotes graft epithelial repair. Unlike *Hif1α* ablation, 2-DG treatment did not impact IL-1β^+^ Mo-AM accumulation (**Figure 7F**), although it did reduce the abundance of IL-1β^+^ CD11c^+^ cells (**Figure 7G**). Additionally, when compared to vehicle treatment, 2-DG administration lowered plasma lactate levels in Atg16l1-deficient lung recipients (**Figure 7H**).

**Figure 7.**
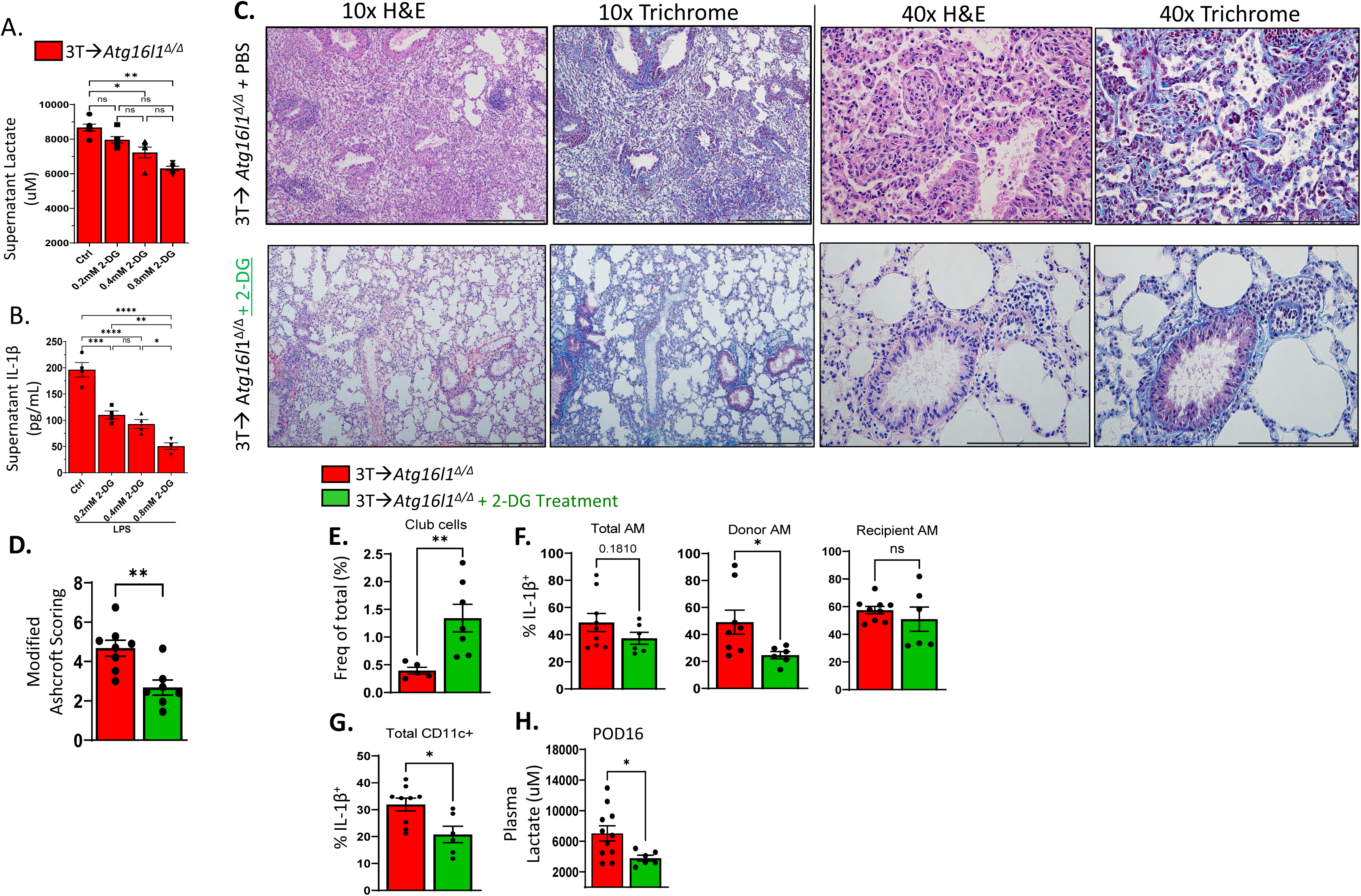
Pharmacological blockade of glycolysis in *Atg16l1^Δ/Δ^* lung recipients prevents CLAD Dose-dependent effects of 2-DG on (**A**) lactate and (**B**) IL-1β production in LPS-stimulated *Atg16l1^Δ/Δ^* BMMs. *Atg16l1^Δ/Δ^* lung recipients of 3T FVB lungs received a daily intraperitoneal injection of PBS or 2-DG (750 mg/kg) from POD 7 to POD 16. POD 16 allograft histology as shown by representative images of (**C**) H&E and trichrome staining and blindly graded for (**D**) modified Ashcroft score. POD 16 allograft cells assessed for (**E**) club cell abundance relative to CD45^-^ cells, (**F**) intracellular AM IL-1β expression and (**G**) intracellular IL-1β expression from CD11c^+^ cells. (**H**) Lung recipient plasma was collected on POD 16 and evaluated for lactate levels. Data in **A-B, D-H** are represented as mean ± SEM. One-way ANOVA (**A-B**), Mann Whitney U T-test (**D-H**). *P < 0.05; **P < 0.01, ***P < 0.001, ****P < 0.0001. Scale bars: 500 μm for 10x, 200 μm for 40x.

## Discussion

We report that lung recipients carrying two *ATG16L1* rs2241880 G alleles significantly increase the risk of accelerated CLAD development after lung transplantation. Notably, the rs2241880 G minor allele frequency (MAF) in our cohort of lung transplant recipients was approximately 70%. Although this appeared surprisingly high, prior work analyzing rs2241880 G allele occurrence in individuals of European ancestry showed a MAF of 47% (*17*). Interestingly, in the same study, rs2241880 AG carriers also had a significant risk for Crohn’s disease. We also observed a shift towards earlier CLAD in recipients carrying rs2241880 AG alleles, although it did not quite reach statistical significance.

The high proportion of rs2241880 GG carriers in lung recipients relative to the general population suggests that there may be a link to lung inflammation and the development of severe lung disease, leading to transplantation in our cohort. To this end, a study performed in Australia revealed that, among smokers, the risk for Crohn’s disease was increased over seven-fold in rs2241880 GG carriers relative to non-smokers (*10*).

Multiple splice variants of *ATG16L1* have been reported in exon 9, which contains the rs2241880 Crohn’s disease susceptibility locus (*17*). Importantly, all of these splice variants contained the threonine to alanine substitution. Recent work on rs2241880 SNPs reveals sharp demographic differences, potentially challenging the generalizability of our findings. Indeed, Murthy and colleagues show that among Northern and Western European ancestries, the MAF is 0.57 with a GG frequency of 0.29, populations of African origin in southwestern regions of the United States have a MAF of 0.26 and a GG frequency of 0.053, people of Mexican ancestry in the Los Angeles area have an MAF of 0.30 and a GG frequency of 0.123. In contrast, people of Chinese descent in Denver, Colorado, have a MAF of 0.236 and a GG frequency of 0.056 (*45*). However, similar to most lung transplant center demographics across the United States (*67*), the majority of lung recipients in our program were of non-Hispanic and European ancestry, which likely contributed to the enrichment of rs2241880 GG carriers. Future studies will be necessary to validate our observations in other cohorts.

The reported role of Atg16l1 in xenophagy suggests that accelerated CLAD development may be due to exacerbated inflammatory responses to infection. Atg16l1 deficiency leads to impaired clearance of intracellular bacteria due to defective autophagosome generation and lysosomal degradation, contributing to enhanced inflammatory signaling in a mouse model of inflammatory bowel disease (*45*). However, in a mouse model of urinary tract infection, Atg16l1 deficiency elevated IL-1β production and increased uptake of *E. coli*, which protected from infection (*65*). In our multivariable model, we accounted for infectious agents known to increase the risk of CLAD, including bacterial, viral, and fungal infections (*27, 49, 50, 53, 54, 62*). Unfortunately, we were unable to determine if infections were more common in rs2241880 GG carriers due to insufficient cohort size. Nevertheless, it is noteworthy that ATG5 plays a similar role in antimicrobial defense; however, carrying ATG5 SNPs did not increase CLAD risk, suggesting that *ATG16L1^T300A/T300A^* promotes CLAD development independently of infection (*13*).

Transcript profiling of Atg16l1-deficient macrophages revealed substantial evidence of mitochondrial stress. There was a marked loss of nuclear encoded transcripts that direct the synthesis of many ETC subunits, with ETC I components being the most affected. Under high mitochondrial membrane potential, ETC I becomes a significant source of superoxide ions, which is thought to be the result of reverse flow of electrons from succinate oxidation at ETC II (*6, 7, 44, 71*). Additionally, ETC III can also generate high amounts of ROS when ETC components are damaged or electron flow is impaired (*16*). Regarding these reports, we noted increased mitochondrial membrane potential, enhanced mitochondrial mass, and elevated superoxide production in Atg16l1-deficient macrophages relative to their wildtype counterparts. We hypothesized that high mitochondrial mass was due to poor clearance of damaged mitochondria through mitophagy. Examining macrophages generated from mice that express the mt-Keima mitophagy flux reporter protein, we detected reduced Atg16l1 T300A-dependent decrement of mitochondrial flux into lysosomes. This observation is consistent with a recent report demonstrating that a myeloid-targeted gene deletion of *Atg16l1* inhibits mitophagy after partial hepatectomy in mice (*72*). Additionally, along with a marked attenuation in CLAD severity, enhanced IL-1β expression was reversed in recipients with Atg16l1-Hif1α doubly deficient macrophages. Although the precise mechanisms are not clear, increased ETC ROS activity is reported to promote Hif1α stabilization (*16*), suggesting that augmented IL-1β expression results from damaged mitochondria accumulation in Atg16l1-deficient macrophages. Future studies focusing on the role of mitochondrial ROS in promoting Hif1α-mediated IL-1β production and its potential impact on CLAD pathogenesis are warranted.

Besides promoting IL-1β expression, Hif1α stabilization enhances glycolysis (*66*).

Measurements of the glycolysis-related transcripts, extracellular acidification rate, lactate production, and glycolytic flux all indicated that Atg16l1 is a critical regulator of glycolysis in macrophages. Surprisingly, only the deletion of the *Hif1α* allele from Atg16l1^ΔΔ^ BMM was sufficient to reduce glycolytic activity, IL-1β expression, and significantly attenuate CLAD severity in *Atg16l1^ΔΔ^* recipients. These data suggest a critical role of Hif1α stabilization in allograft injury. Several earlier observations may explain this observation by relating mitochondrial damage to altered metabolism. First, LPS, in addition to generating mitochondrial ROS, could further promote Hif1α stability through its ability to raise succinate levels (*59, 66*). Succinate directly inhibits the activity of HIF-α-prolyl hydroxylase, an enzyme that targets Hif1α for destruction (*9*). Second, the buildup of damaged mitochondria in Ag16l1-deficient macrophages may be further exacerbated by LPS-mediated damage to ETC components (*3*), increased mitochondrial membrane potential (*42*), and mitophagy inhibition in the presence of IFN-γ (*52*), a cytokine that is produced by macrophages following LPS stimulation and is elevated in human lung transplant airspaces (*33*).

Augmented glycolysis has been linked to lung disease. For example, patients with idiopathic pulmonary fibrosis and acute lung injury have increased circulating lactate levels (*25, 70, 73*). In a murine model of silicosis-induced pulmonary fibrosis, 2-DG treatment inhibited Hif1α stabilization, reduced the accumulation of glycolytic transcripts, and prevented the development of fibrotic lesions (*39*). However, the role of glycolysis in allograft tolerance is less understood. In mice, simultaneously targeting several facets of glucose metabolism using a combination of 2-DG, metformin, and 6-dao-5-oxo-L-norleucine, the latter of which inhibits gluconeogenesis, prolonged both skin and heart allograft survival (*31*). Using 2-DG treatment alone, we were able to attenuate CLAD severity in *Atg16l1^ΔΔ^* recipients. Interestingly, reduced allograft injury could be potentially explained by previous observations that 2-DG interferes with protein glycosylation, leading to endoplasmic reticulum stress and potential toxicity (*61*). Nonetheless, systemic administration is reported to be safe and effective in several clinical trials, including the sensitization of human osteosarcoma and non-small cell lung cancers to chemotherapy, as well as in the treatment of COVID-19 (*2, 41, 48, 63*).

There are several limitations to our study. This includes the lack of a validation cohort and the absence of SNP data on donors. Additionally, the use of a mouse transplant model may not fully represent the spectrum of CLAD pathology, given the phylogenetic differences in club cell distribution and lung airway anatomy with respect to humans. Still, we have identified a highly prevalent ATG16L1 hypomorphic mutation enriched in human lung recipients that is linked to accelerated CLAD development.

When modeled in mouse lung transplants, this mutation is found to prevent allograft tolerance through Hif1α-dependent pathways that drive glycolysis and inflammation. Thus, our findings suggest the potential of immunometabolism-based therapies for CLAD, with implications for personalized medicine.

## Methods

### CLAD Diagnosis

This study was approved by the Institutional Review Board at Washington University in Saint Louis. All patients provided consent for inclusion in this study were included in a study database ranging from 1992 to 2013. Lung transplant recipients who underwent lung transplantation at Washington University in Saint Louis (n=560) were sequentially enrolled starting December 2009. Clinical data from recipients, donors, perioperative, rejection, and infection data were collected through chart review and entered into a REDCap database. Peak Forced Expiratory Volume (FEV1) in liters per second was calculated from the average of the two highest FEV1 measures. The percentage of the peak FEV1 for every subsequent measurement was calculated. The date of the first FEV1 that was <90% of the peak FEV1 was defined as the onset of CLAD in accordance with recommendations of the International Society for Heart and Lung Transplant (*69*). Sequencing of ATG16l1 and ATG5 SNPs was performed by DNA Genotek (Kanata, Ottawa, Canada).

### Mice and Lung Transplant CLAD Model

Animal studies were approved by the Institutional Animal Care and Use Committee at Washington University in Saint Louis. 3T mice on the FVB background have been previously described and were used as donors for all studies (*37, 38*). *Lyz2^Cre^* and *Hif1α^flox/flox^* mice were purchased from Jackson Laboratory. *Atg16l1^fl/fl^* were a gift from Dr. Skip Virgin of Washington University of St. Louis. *Atg16l1^T316A/T316A^* mice were a gift from Genentech (San Francisco, CA). Mt-Keima mice were generated by the targeted injection of mt-Kiema cDNA with a CMV promoter into B6/CBA Rosa Talens hybrid nuclei. Founder mt-kiema mice were selected by screening for the brightest population of peripheral blood leukocyte fluorescence as observed through a 620/20 nm bandpass filter following 488 nm laser excitation. The founders were then backcrossed at least eight times onto the C57Bl/6 background, guided by QTL analysis performed by Jackson Laboratories. The mouse left orthotopic lung transplantation procedure have been described previously (*47*). Recipients were given 250 μg of CD40L Ab (clone MR1) on POD0 and 200 ug of mouse recombinant CTLA4 Ig on POD2 intraperitoneally to induce tolerance (*37*). To induce moderate club cell injury, Doxycycline was ingested via food (625 mg/kg; ENVIGO) and water (2 mg/ml, MilliporeSigma) for 18 hours instead of 2.5 days (*37*). Mice lung recipients were euthanized on POD16 and lungs processed for histological and flow cytometric analysis.

### Hydroxyproline Assay

A Hydroxyproline Assay Kit (Catalog No.: MAK008) was used to measure the transplanted lung graft collagen in accordance with the manufacturer’s instructions. Briefly, homogenized lung tissue was hydrolyzed with 12 M of HCl at 120 °C for 3 hours. Supernatants were added to the plate and dried in a 60 °C oven overnight. The hydroxyproline concentration was determined by the reaction of oxidized hydroxyproline with 4-(dimethylamino) benzaldehyde, which generates a colorimetric product and is measurable by absorbance at 560 nm with a BioTek Synergy HTX plate reader.

### Histological Analysis and Ashcroft Scoring

Mouse lungs were processed in buffered formalin, paraffin-embedded, and stained with Hematoxylin and Eosin or Gomori trichrome. Lung section slides were scanned with a Zeiss Axio Scan 7 Brightfield Fluorescence Scanner with 20-fold magnification.

Scanned images were divided into 1200 x 1200 μm tiles using QuPath-0.4.3 software. Tiles with dominant bronchial or vascular structures were excluded from scoring. Ten tiles were randomly selected and scored blindly by two experienced researchers according to the definition of the Modified Ashcroft Scale (*19*). In accordance with this method, a predominant degree of fibrosis of each tile was assessed on a scale from 0 to 8. The scales were summarized and divided by the number of tiles to obtain a fibrotic index for each transplanted lung.

### Bulk RNASeq Analysis

Lungs from Atg6l1-deficient (*Atg16l1^ΔΔ^*) and *Atg16l1^flox/flox^* control mice were FACS sorted for live cells (Zombie^-^), then recipient-derived alveolar macrophages (AM; CD45.2^+^, CD11c^+^, SiglecF^+^) and interstitial macrophages (IM; CD45.2^+^, CD11c^+^, CD11b^+^, CD64^+^, SiglecF^-^). These were directly sorted into Trizol (ThermoFisher) for immediate processing by the Washington University School of Medicine Genome Technology Access Center Core Facility (Saint Louis, MO) for Bulk RNA sequencing on Illumina NovaSeq 6000 sequencing system. For bulk-RNASeq of BMM and monocytes were isolated from bone-marrow with a mouse monocyte isolation kit (Miltenyi), cultured for in 15 ng/mL GM-CSF for 7 days then stimulated with E.Coli derived LPS (Strain OB111, Sigma Aldrich) at 100 ng/mL overnight before collecting for sequencing.

### Macrophage Culture and Analysis

Bone marrow monocytes isolated with a Monocyte isolation kit (Miltenyi) in accordance with manufacturers recommendations. Monocytes were differentiated into BMM for 7 days in RPMI 1640 media (ThermoFisher) with 10% Fetal Bovine Serum with 15 ng/mL of recombinant mouse GM-CSF (R&D). BMM overnight stimulation with either phosphate buffered saline (PBS) control or E. Coli LPS (Sigma; Strain O111:B4) for 18 hours. Supernatant lactate production was measured using an EtonBio Lactate Assay kit (San Diego, CA). For glycolysis inhibition, cells were washed and treated with 2-Deoxy-D-glucose (0, 0.2mM, 0.4mM, 0.8mM) for 4 hours before overnight treatment of either PBS or LPS. For plasma lactate measurements, blood was collected from mouse lung transplant recipients on postoperative day (POD) 16 at sacrifice. Blood was centrifuged at RT for 20min at 5000rpm and plasma stored at-80^0^C until lactate measures. Unpaired t-test with nonparametric correction or ANOVA were used when appropriate.

### Mitochondria and Extracellular Flux Analysis

Glycolytic activity was measured using the Seahorse platform (Agilent SXF96, Wave 2.3 version). 100,000 BMM were seeded per well in a 96-well Seahorse plate and stimulated with 100 ng/mL LPS. Macrophages were incubated with Seahorse DMEM media supplemented with 10 mM glucose, 2 mM L-Glutamax, 1 mM Sodium Pyruvate at a pH 7.4 and incubated at 37^0^C without CO_2_ for 1 hr for the mitochondrial stress test.

Cultures received sequential treatment with 1 μM Oligomycin, 1.5 μM Carbonyl cyanide p-trifluoromethoxyphenylhydrazone (FCCP) and a 1:1 mixture of 1 μM Rotenone & Antimycin spaced at approximately 27-minute intervals to determine continual oxygen consumption rate (OCR) changes. For the glycolysis stress test macrophages were washed and replaced with glucose-free Seahorse medium supplemented with 2 mM glutamate, incubated at 37 °C without CO_2_ and received sequential delivery of 10mM glucose, 1 μM Oligomycin and 50 mM 2-DG spaced at 27-minute intervals to measure continuous alterations in extracellular acidification rate (ECAR).

### Radiometric Glycolytic Flux Assay

Glycolytic flux was determined by measuring [5-^3^H]-glucose metabolism (*34, 35*). Tritium-labeled D-[5-^3^H(N)]-glucose tracer (Revity) was added at 0.4 μCi/mL to BMM after LPS stimulation and incubated for 2 hours at 37°C to allow for glycolysis. The supernatant was transferred to an Eppendorf tube containing 50 μL of 5N HCl. The microcentrifuge tubes were then placed in 20 ml scintillation vials containing an equivalent volume of water for humidification. The vials were then capped with a piece of hanging Whatman filter paper and sealed for three days at room temperature. ^3^H_2_O, generated by glycolysis, collects on the hanging filter paper through evaporative diffusion. This filter is then placed in scintillation fluid to measure ^3^H_2_O activity. Cell-free samples containing 1 μCi of ^3^H-glucose were used as unmetabolized controls. Cells were lysed for protein measurements with the Biuret reagent assay (Millipore Sigma) to correct for cell mass.

### Semi-quantitative real-time PCR

RNA was extracted from bone marrow-derived macrophages using the RNeasy Mini Kit (QIAGEN), and reverse transcribed with iScript Reverse Transcription Supermix (BioRad) according to manufacturer’s instructions. Transcripts of indicated genes were assessed using TaqMan Gene Expression Assays (ThermoFisher) and normalization against housekeeping gene β-actin.

### Immunohistochemical Staining

Monocytes were plated on sterile glass coverslips in a 12-well plate. After differentiation into macrophages, the cells were washed with cold PBS three times, fixed with 4% paraformaldehyde in PBS pH 7.4 for 20 min at room temperature and permeabilized with 0.1% Triton X-100 for 20 min at room temperature. The cells were blocked with 2% BSA in PBS for 30 min and incubated with ATG16L1(clone D6D5; Cell Signaling) and Tom20 antibodies (clone 4F3; Abnova) at a 1:500 dilutions overnight at 4°C. Cells were washed for three times in PBS, incubated with Goat anti-rabbit-IgG-PE (ThermoFisher, cat # P-2771MP) and Goat anti-mouse IgG-FITC (ThermoFisher Cat # 31569) secondary antibodies at a 1:1000 dilutions and DAPI in 2% BSA for one hour at 4°C. The coverslips were then mounted with 7μl mounting medium and sealed with nail polish prevent drying and movement under microscope.

### Mitophagy Analysis

Monocytes were purified from the bone marrow of C57/BL6 *(B6)^mt-Keima/+^* and *B6 ^mtKeima/+^Atg16l1^T316A/T316A^* mice, then seeded on a 35-mm glass chamber dish containing a 1.5 mm coverslip (MatTek) at a density of 3 × 10^5^ cells per well in 2 ml of RPMI140 supplemented with 10% Fetal Bovine serum. Monocytes were incubated overnight at 37°C in 5% CO_2_. Images were generated by excitation at 458 nm (cytoplasmic) and 561 nm (lysosomal) using a Zeiss LSM 780 confocal microscope (Carl Zeiss MicroImaging). Levels of mitophagic flux were assessed by calculating the ratio of cytoplasmic to lysosomal-induced fluorescence intensity. BMM were stained with MitoSox Red (ThermoFisher), MitoTracker Green (Thermofisher) and MitoTracker Red (Thermofisher) to assess mitochondrial reactive oxygen production, mass, and membrane potential, respectively, in accordance with the manufacturer’s recommendations.

### Flow Cytometric Analysis

Lung tissue was digested and prepared for single-cell suspension as previously described (*47*). Live/dead cell staining was performed with the Zombie Violet Fixable Viability Kit (Biolegend). Cell surface staining was conducted with the following antibodies: CD45 (30-F11, eBioscience), CD45.2 (104, Biolegend), CD45.1 (A20, Biolegend), SiglecF (E50-2440, BD Bioscience), CD64 (X54-5/7.1, eBioscience), CD11c (N418, Biolegend), CD31 (390, Biolegend), CD11b (M1/70, Biolegend), CD34 (HM34, Biolegend), and CD326 (G8.8, Biolegend), H-2Kq (KM114, Biolegend), Ki-67 (16A8, Biolegend), and polyclonal rabbit anti-mouse CCSP (Seven Hills Bioreagents, Cat# WRAB-3950) were conducted with an intranuclear Transcription Factor Staining Buffer Kit (Invitrogen) in accordance with the manufacturer’s recommendations. For intracellular IL-1β staining, lung single-cell suspensions were treated with Golgi plug (BD Biosciences) and stained with IL-1β antibodies (B122; BioLegend) with a Cytofix/Cytoperm kit (BD Biosciences) in accordance with the manufacturer’s recommendations.

## Statistical Analysis

Univariate and multivariate analyses were conducted to adjust for contributing factors. Log-rank analysis was performed to assess freedom from CLAD and the probability of survival with pairwise comparisons. For comparisons between three groups, ANOVA was used with a Tukey’s HSD post-hoc test, and for non-pairwise comparisons, the Mann-Whitney U non-parametric Test was employed where indicated.

## List of Supplementary Data

Table S1. Data retrieved from the chart review for analysis. Table S2. Infections retrieved from chart review for analysis

Fig S1. Lung recipients who carry Atg5 SNPs do not have reduced freedom from CLAD or allograft survival

Fig S2. CLAD development in *Atg16l1^T316A/T316A^* lung transplant recipients.

Fig S3. Increased glycolysis in *Atg16l1^T316A/T316A^* BMM.

Fig S4. Upregulation of Hif1α glycolytic and inflammatory target expression

Fig S5. Ablation of both *Hif1α* alleles from Atg16l1-deficient BMM attenuates glycolysis and IL1β expression

Fig S6. *Hif1a^Δ/+^Atg16l1^ΔΔ^* lung recipients have less evidence of CLAD relative to *Atg16l1^ΔΔ^* lung recipients.

Fig S7. The ablation of one *Hif1α* allele from Atg16l1 BMM is sufficient to reduce lactate and IL-1β production.

Movie S1. Colocalization of Atg16l1 and mitochondria in BMMs

## Funding

National Institutes of Health T32HL007317 (MC)

National Institutes of Health, NHLBI PRIDE HL126140 (MC) American Thoracic Research Grant (MC)

National Clinical and Translational Science Awards KL2TR002346 (MC) National Institutes of Health, R01HL094601 (AEG, DK)

National Institutes of Health, P01AI116501 (DK, ASK, AEG) National Institutes of Health, R01 HL167277 (AEG) National Institutes of Health, R01 HL169189 (AEG)

## Author contributions

Conceptualization: MC, FL, AEG

Methodology: MC, FL, DZ, CC, ZL, JZ, RE, VC, KP, YT, AEG Investigation: MC, FL, DZ, CC, ZL, JZ, CB, RE, VC, KP, YT, AEG

Visualization: MC, FL, DZ, AEG

Funding acquisition: MC, DK, ASK, AEG, Project administration: MC, AEG Supervision: LT, DB, RH, SLB, DK, AEG Writing – original draft: MC, ZL, AEG

Writing – review & editing: MC, LT, DB, RH, SB, DK, AEG

We thank the Genome Technology Access Center in the Department of Genetics at Washington University School of Medicine for help with genomic analysis. The Center is partially supported by NCI Cancer Center Support Grant #P30 CA91842 to the Siteman Cancer Center and by ICTS/CTSA Grant# UL1TR002345 from the National Center for Research Resources (NCRR), a component of the National Institutes of Health (NIH), and NIH Roadmap for Medical Research. This publication is solely the responsibility of the authors and does not necessarily represent the official view of NCRR or NIH.

