## Supplemental Data for "Atg16l1 promotes lung transplant tolerance by regulating glycolysis in macrophages"

| Recipient Data | Donor Data | Perioperative Data | Rejection Data | Infection Data |
|---|---|---|---|---|
| Gender<br>Race<br>Ethnicity<br>Age at transplant<br>ABO group<br>RhD antigen<br>Height at transplant<br>Weight at transplant<br>BMI at transplant<br>Mean pulmonary artery pressure<br>Presence of pulmonary hypertension<br>Primary diagnosis<br>Lung allocation score<br>Date of expiration or retransplantation<br>Time of graft survival<br>CMV serology | Gender<br>Race<br>Ethnicity<br>Age<br>ABO group<br>RhD antigen<br>Non-heart beating donor<br>Cause of death<br>Smoking history<br>$PaO_2$<br>$FiO_2$<br>Abnormal chest x-ray<br>Abnormal bronchoscopy<br>Extended criteria<br>CMV serology | Date of lung transplant<br>Transplant type<br>Maximum ischemia time<br>Cardiopulmonary bypass use<br>Cardiopulmonary bypass time<br>PGD grade 0-6 hours<br>PGD grade 24 hours<br>PGD grade 48 hours<br>PGD grade 72 hours | Cumulative A rejection score at 1, 3, 6, 12, and 24 months<br>Cumulative B rejection score at 1, 3, 6, 12, and 24 months<br>Number of biopsies performed by 1, 3, 6, 12, and 24 months<br>Time to A1 and A2<br>Time to B1 and B2 | Species of first *Aspergillus* infection post-transplant<br>Time to first *Aspergillus* infection<br>Species of first Gram-positive infection post-transplant<br>Time to first Gram-positive infection<br>Species of first Gram-negative infection post-transplant<br>Time to first Gram-negative infection<br>Species of first CARV infection post-transplant<br>Time to first CARV infection<br>Location of first CMV infection post-transplant<br>Time to first CMV infection |

**Supplementary Table 1. Data retrieved from chart review for analysis.**

| **Infectious Agents, n=229** | **n (%)** |
| --- | --- |
| Gram-positive bacteria | 63 (27.51) |
| *Staphylococcus aureus* | 46 (20.09) |
| Other | 17 (7.42) |
| Gram-negative bacteria | 119 (51.97) |
| *Pseudomonas aeruginosa* | 53 (23.14) |
| *Stenotrophomonas maltophilia* | 50 (21.83) |
| Other | 6 (2.62) |
| Cytomegalovirus | 164 (71.62) |
| Blood | 137 (59.83) |
| Airway | 18 (7.86) |
| Blood and airway | 1 (0.44) |
| Community-acquired respiratory virus | 82 (35.8) |
| Influenza | 5 (2.18) |
| Respiratory syncytial virus | 15 (6.55) |
| Adenovirus | 2 (0.87) |
| Other | 60 (26.20) |
| *Aspergillus* | 83 (36.24) |
| *Aspergillus fumigatus* | 22 (9.61) |
| *Aspergillus niger* | 15 (6.55) |
| *Aspergillus flavus* | 19 (8.30) |
| Other single species | 20 (8.73) |
| Multiple single species | 7 (3.06) |

**Supplementary Table 2. Infections retrieved from chart review for analysis.**

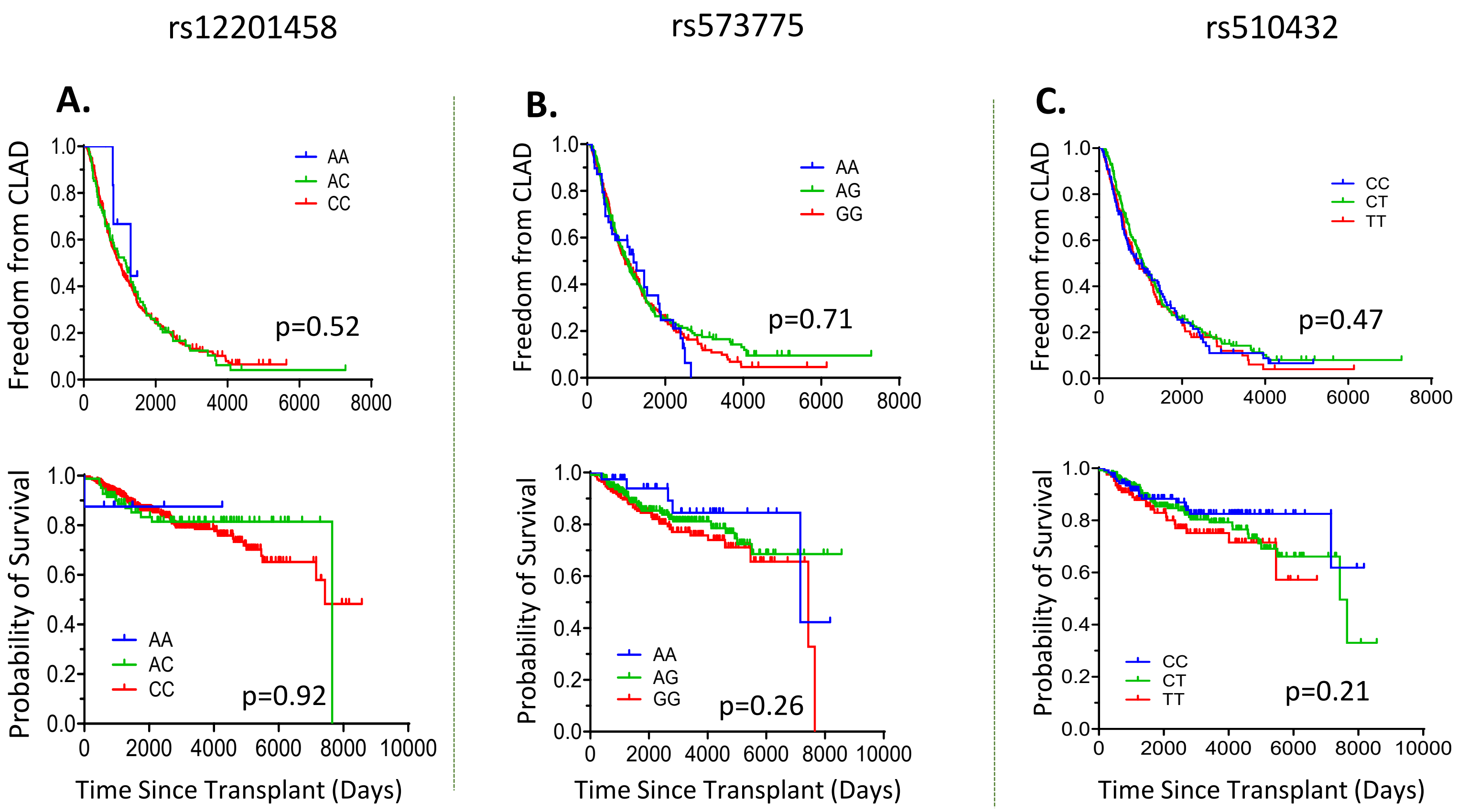


**Supplementary Figure 1. Lung recipients who carry *ATG5* SNPs do not have reduced freedom from CLAD or allograft survival**

The percentage of recipients free of CLAD (upper panel) and their probability of transplant survival (lower panel) with respect to carrying Atg5 variant (**A**) rs12201458, (**B**) rs573775, and (**C**) rs510432 when stratified by genotype.

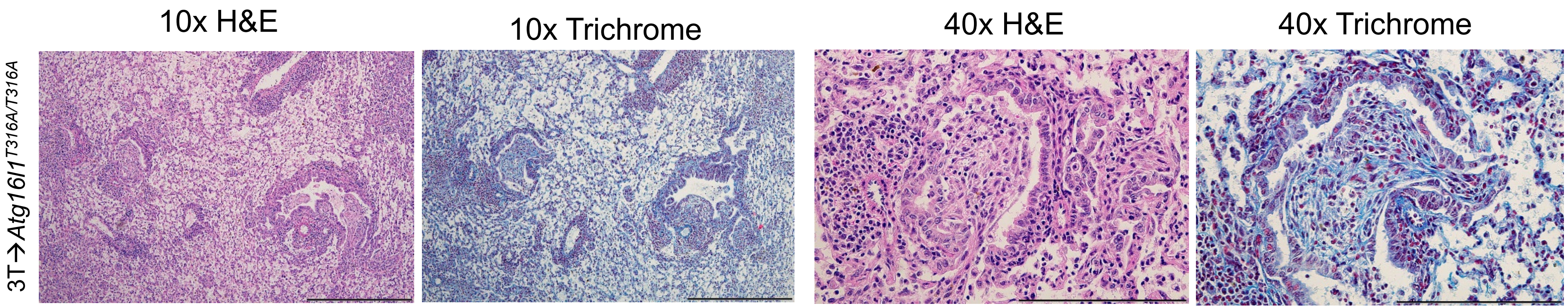


**Supplementary Figure 2. CLAD development in *Atg16l1*$^{T316A/T316A}$ lung transplant recipients.**

Representative H&E and Gomori trichrome staining of *Atg16l1*$^{T316A/T316A}$ and B6 wildtype lung allograft tissue (n≥6/group). Scale bars: 500 μm for 10x, 200 μm for 40x magnification.

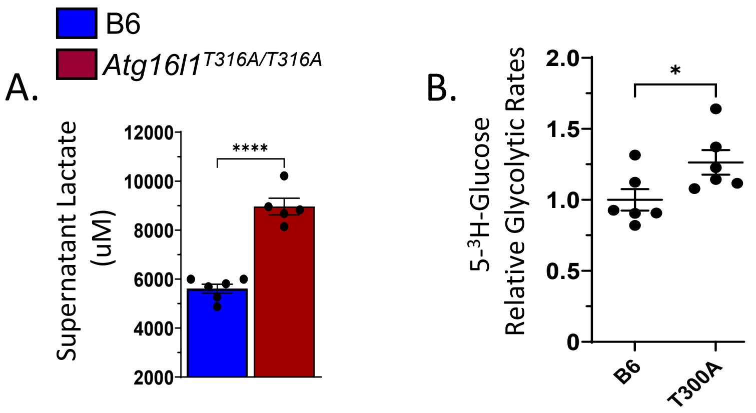


**Supplementary Figure 3. Increased glycolysis in *Atg16l1$^{T316A/T316A}$* BMM.**

BMMs from B6 or *Atg16l1$^{T316A/316A}$* mice were evaluated for (**A**) supernatant lactate levels and (**B**) glycolytic flux of 5-$^{3}$H-glucose. Data are represented as mean ± SEM. A two-sided Mann-Whitney U test was conducted for (**A-B**). n≥5/group *P < 0.05; ****P < 0.0001.

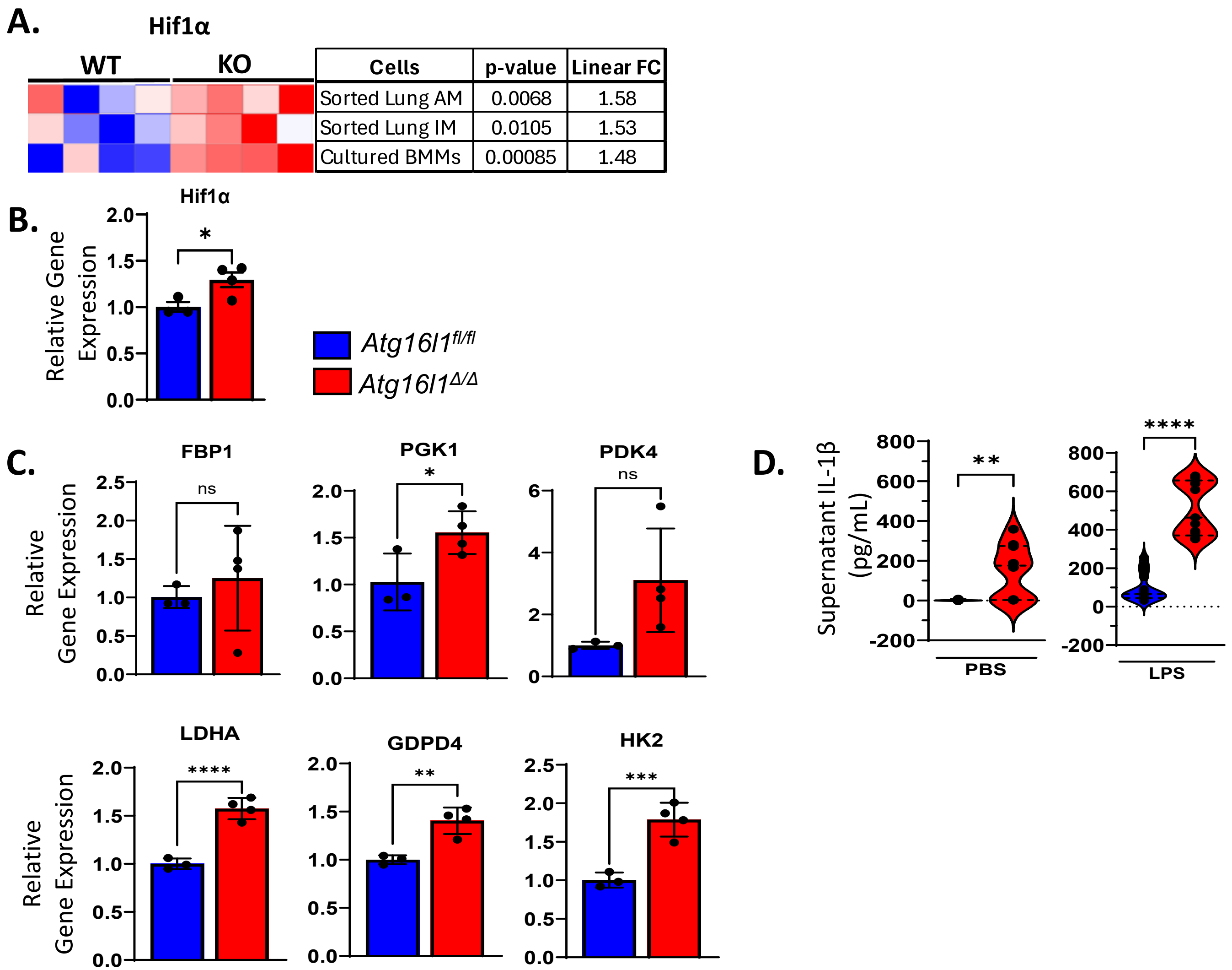


**Supplementary Figure 4. Upregulation of Hif1α glycolytic and inflammatory target expression**

(**A**) Heat map of *Hif1α* transcripts in control *Atg16l1*$^{fl/fl}$ and *Atg16l1*$^{\Delta/\Delta}$ FACS-sorted lung alveolar macrophages (AM), interstitial macrophages (IM), and BMMs. (**B**) Transcript accumulation of *Hif1α* was normalized to the β-Actin mRNA levels in *Atg16l1*$^{fl/fl}$ and *Atg16l1*$^{\Delta/\Delta}$ BMMs (n≥3/group). (**C**) Glycolysis-related transcript levels normalized to β-Actin mRNA in control *Atg16l1*$^{fl/fl}$ or *Atg16l1*$^{\Delta/\Delta}$ BMMs (n≥3/group). (**D**) *Atg16l1*$^{fl/fl}$ and *Atg16l1*$^{\Delta/\Delta}$ BMM spontaneous and LPS-induced IL-1β expression. Data are represented as mean ± SEM. Two-sided Mann-Whitney U test (**B-D**). *$P < 0.05$; **$P < 0.01$, ****$P < 0.0001$.

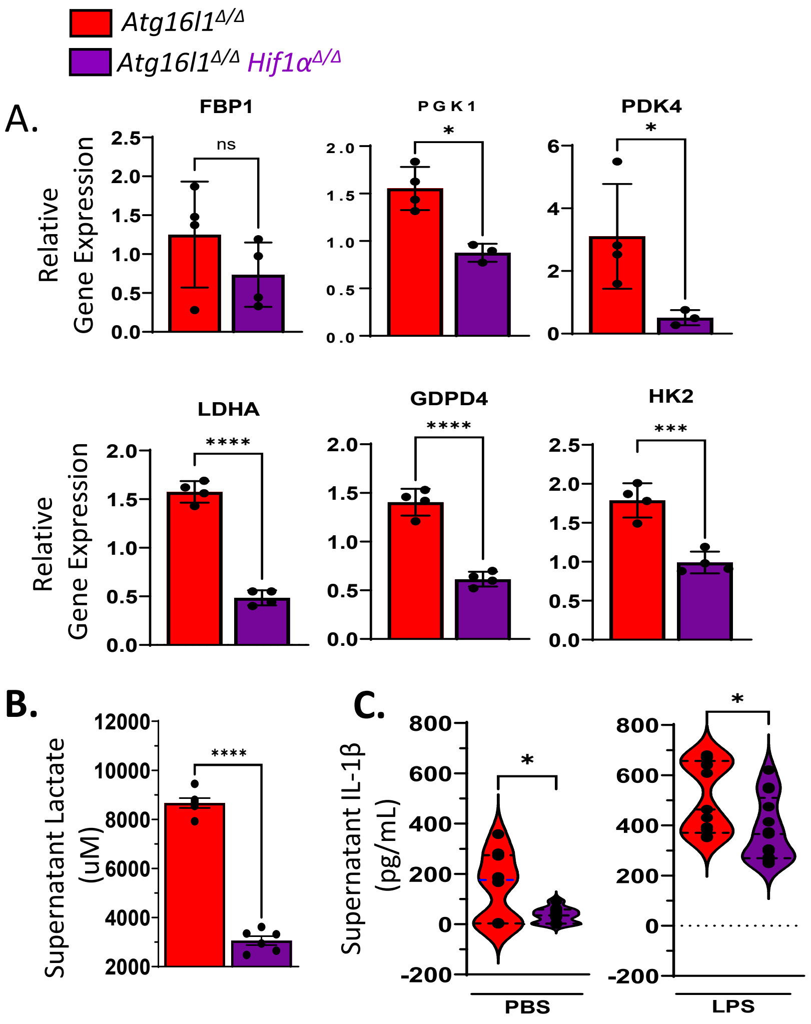


**Supplementary Figure 5. Ablation of both *Hif1α* alleles from Atg16l1-deficient BMM attenuates glycolysis and IL1β expression**

(**A**) Accumulation of glycolysis-related transcripts normalized to *β-Actin* (n=4/group). (**B**) Supernatant lactate levels (n≥4/group), and (**C**) IL-1β accumulation following overnight 100ng/ml LPS stimulation of *Atg16l1*$^{\Delta/\Delta}$ and *Atg16l1*$^{\Delta/\Delta}$ *Hif1α*$^{\Delta/\Delta}$ BMMs. Data are represented as a mean ± SEM. Two-sided Mann-Whitney U test (**A-C**). *P < 0.05; **P < 0.01, ***P < 0.001, ****P < 0.0001.

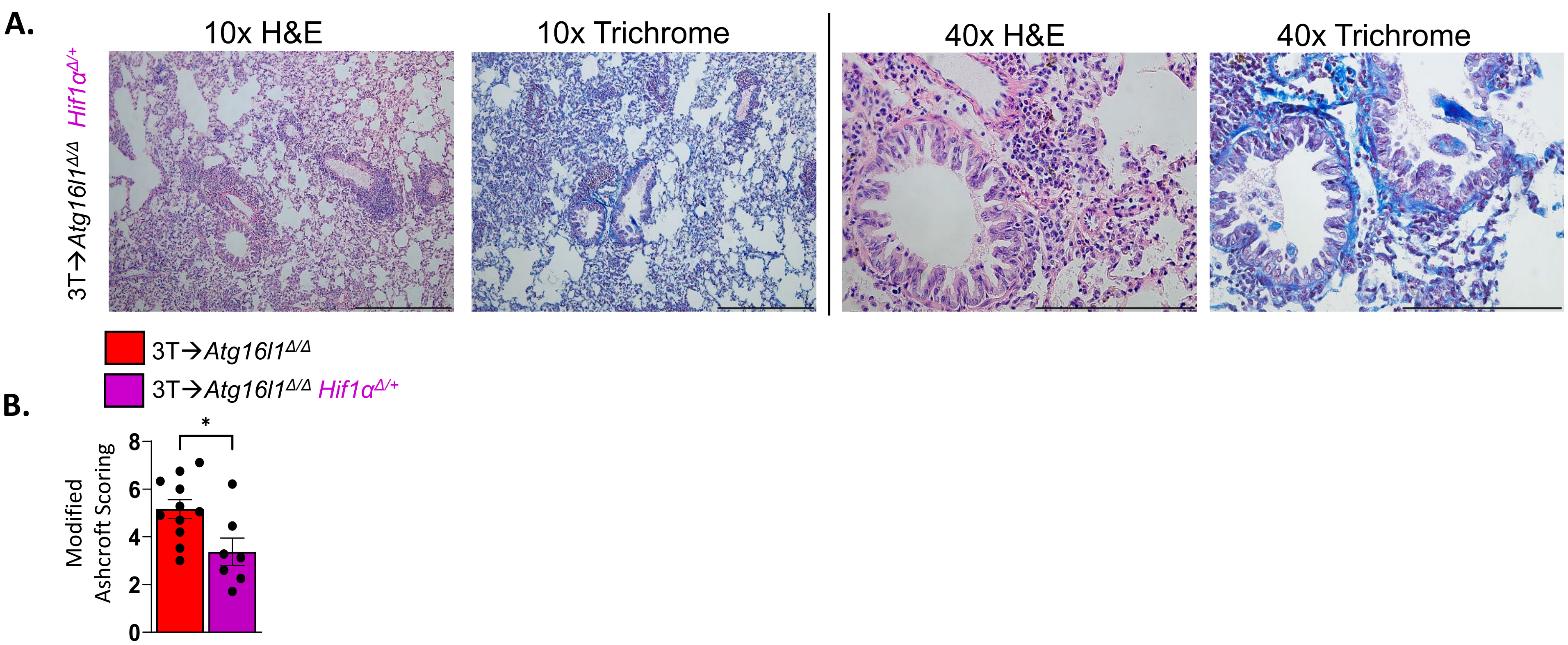


**Supplementary Figure 6.** ***Hif1α***$^{\Delta/+}$ ***Atg16l1***$^{\Delta/\Delta}$ **lung recipients have less evidence of CLAD relative to** ***Atg16l1***$^{\Delta/\Delta}$ **lung recipients.**

Representative **(A)** H&E and trichrome staining and **(B)** modified Ashcroft scoring of allograft tissue from *Atg16l1*$^{\Delta/\Delta}$ and *Atg16l1*$^{\Delta/\Delta}$ *Hif1α*$^{\Delta/+}$ transplant recipients (n≥6/group).

Data in (**B**) are represented as mean ± SEM. Mann-Whitney U t-test. *$P < 0.05$.

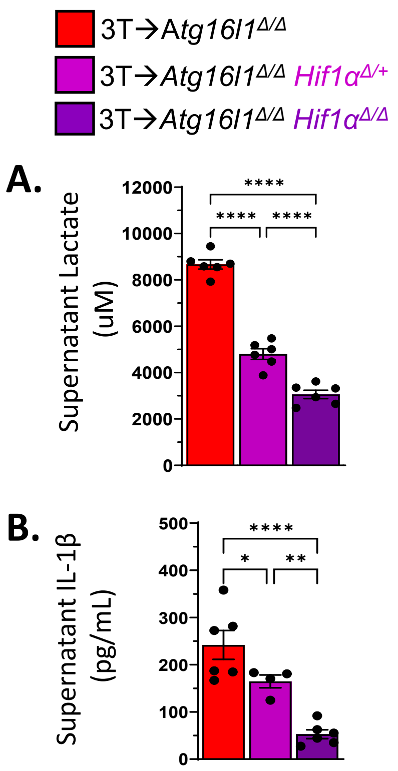


**Supplementary Figure 7. The ablation of one Hif1α allele from Atg16l1 BMM is sufficient to reduce lactate and IL-1β production.**

BMMs from *Atg16l1*$^{\Delta/\Delta}$*, Atg16l1*$^{\Delta/\Delta}$ *Hif1α*$^{\Delta/+}$, and *Atg16l1*$^{\Delta/\Delta}$ *Hif1α*$^{\Delta/\Delta}$ mice were evaluated for supernatant (**A)** lactate and (**B**) IL-1β levels. Data are represented as mean ± SEM. Two-sided Mann-Whitney U test (**A-B**) for n≥5/group where *P < 0.05; **P < 0.01, ****P < 0.0001.

### 3D View of Co-localization between ATG16L1 and Mitochondria

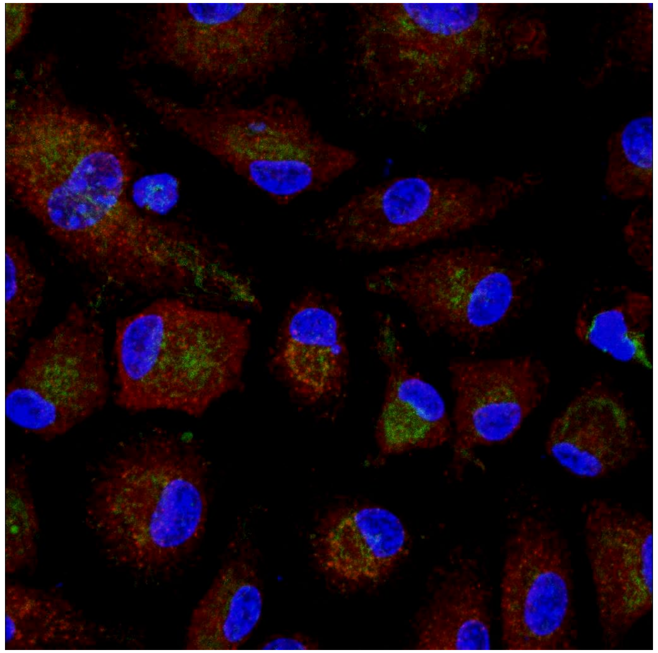

**Supplementary Movie 1. Colocalization of Atg16l1 and mitochondria in BMMs**

Confocal imaging movie of the colocalization of Atg16l1 and mitochondria in BMMs as representative for mitochondria transmembrane protein Tom20 and Atg16l1 co-staining.
